# CA19-9 induces microenvironment remodeling in pancreatic ductal adenocarcinoma

**DOI:** 10.64898/2026.05.22.727290

**Authors:** Jasper Hsu, Hyemin Song, Satoshi Ogawa, Casie S. Kubota, Kristina L. Peck, Emily Jacobs, Lizmarie Garcia-Rivera, Jonathan Zhu, Yuan Sui, Woncheol Jung, Yang Dai, Jan Lumibao, Chelsea R. Bottomley, Kassidy Curtis, Mars Bau, Ellyse Ku, Kahing Kuo, Araceli Herrera Morales, McKenna Stamp, Angelica Rock, Shira R. Okhovat, Tony Hunter, Michael Downes, Ronald Evans, Jingjing Zou, Tae Gyu Oh, Ye Zheng, Andrew M. Lowy, Herve Tiriac, Susan M. Kaech, Dannielle D. Engle

## Abstract

Durable therapeutic efficacy remains a major barrier to improving outcomes for patients with pancreatic ductal adenocarcinoma (PDAC). An immunosuppressive tumor microenvironment (TME) is a hallmark of PDAC and has been demonstrated to be a dominant driver of therapeutic resistance. The aberrant glycan CA19-9 is prevalent in PDAC and drives tumor progression, but the paracrine mechanisms by which it contributes to TME remodeling are unknown. To address this, we mapped TME changes and performed functional analyses using a genetically engineered mouse model (GEMM) harboring Kras^G12D^ mutation and inducible CA19-9 expression. Elevation of CA19-9 led to expansion of antigen-presenting cancer associated fibroblasts (apCAFs) and regulatory T cells (Tregs), which can drive immunosuppression. Antibody blockade of CA19 -9 resulted in significant restoration of normal histology and decreased apCAF and Treg populations. We dissected the paracrine signaling mechanisms that drive this TME remodeling *in vitro* using mouse and human organoid mono- and co-culture models as well as *in vivo* using GEMMs and syngeneic orthotopic transplantation models. CA19-9 induced IL1a and TGFb expression, reprogramming pancreatic mesothelial cells into apCAFs *in vitro*, which in turn directly ligated naïve Cd4^+^ T cells resulting in Treg differentiation in co-cultures. Antibody blockade of IL1a and TGFb in mice led to reduced apCAF and Treg differentiation. We previously reported that CA19-9 modification of the secreted Fbln3 protein increased Egfr engagement and now find that the induction of IL1a and TGFb expression by CA19-9 is dependent on Fbln3 hyperactivation of EGFR signaling. Genetic depletion of Fbln3 led to reduced tumor progression and increased Cd8^+^ T cell infiltration in mice. Together these findings identify a previously unknown signaling axis driving immunosuppressive phenotypes in PDAC, uncovering multiple potential nodes to relieve the immunosuppressive pressures within the PDAC TME.

## INTRODUCTION

Pancreatic ductal adenocarcinoma (**PDAC**) remains one of the most lethal common malignancies and is projected to soon become the 2^nd^ leading cause of cancer death^1^. 10-15% of patients are eligible for surgery, the only cure for PDAC. The standard of care for advanced disease remains chemotherapy combinations, which extend survival by 2-4 months^2–4^. A pro-tumorigenic, immunosuppressive tumor microenvironment (**TME**) is a PDAC hallmark and major contributor to therapeutic resistance^5–8^. Within the TME, cancer associated fibroblasts (**CAFs**) contribute to treatment resistance and immunosuppression^5,6,9,10^. Amongst the first CAF subtypes to be described, inflammatory CAFs (**iCAFs**), distal from tumor cells, express inflammatory factors such as IL6 and LIF. Myofibroblastic CAFs (**myCAFs**) typically reside adjacent to tumor cells and express myofibroblast markers such as aSMA. Since the initial description of these CAFs, additional subtypes have been characterized. Antigen-presenting CAFs (**apCAFs**) express MHC class II and interact with Cd4^+^ T cells, inducing regulatory T cell (**Treg**) differentiation and immunosuppression in mice^11–13^. Naive Cd4^+^ T cells bind to apCAFs in an antigen-dependent manner, resulting in Treg differentiation^13^. In some cancers, depleting Tregs is sufficient to increase Cd8^+^ T cell infiltration and subsequent effectiveness of immune checkpoint therapies (i.e. anti-PD1 and anti-CTLA4)^14–16^. While Tregs are a major immunosuppressive force, there are conflicting reports of their role in PDA^17,18^. Long term genetic elimination of Tregs results in pancreatitis, which is known to accelerate pancreatic tumorigenesis, complicating the interpretation of these depletion studies. Tumor cell-derived IL1a and TGFb induce mesothelial cell differentiation into apCAFs^13^, but also induce opposing myCAF/iCAF differentiation programs. IL1/JAK-STAT3 signaling induces iCAF generation and TGFb induces myCAF differentiation^19^.

Glycosylation is one of the most common and complex types of post-translational modifications. Glycans contribute a significant amount of mass and structural variation, and mostly exist as membrane-bound or secreted glycoconjugates^20,21^. Aberrant glycosylation is a hallmark of cancer^22^ and allows neoplastic cells to reacquire developmental attributes such as receptor activation, cell adhesion, and cell motility, promoting growth and invasion^23^. Changes to glycans result in increased tumor proliferation and immune suppression^23–28^. Despite this, the molecular mechanisms by which glycans promote cancer disease progression remain largely unknown. Carbohydrate antigen 19-9 (**CA19-9**) is tetra-saccharide glycan moiety elevated in serum of 10-30% of pancreatitis patients and 70-90% of pancreatic cancer patients^29,30^. Serum levels of CA19-9 are used as a biomarker of PDAC progression. PDAC patients who do not exhibit elevated CA19-9 have significantly improved prognoses independent of disease stage^25,31,32^. Mice lack CA19-9^33^. Our recent studies using Doxycycline (**Dox**) inducible mouse models demonstrated that CA19-9 elevation caused pancreatitis and that CA19-9 modification of the secreted Fibulin 3 (Fbln3, *Efemp1*) protein led to hyperactivation of Egfr^33^. Coupled with *KRAS* mutation, which drives >90% of human PDAC, CA19-9 expression drove transformation and disease progression, reducing median survival^33^. This was accompanied by substantial remodeling of the TME.

Leveraging the CA19-9 inducible, *Kras*-mutant mouse model and a combination of scRNA-seq and histological approaches, we found that CA19-9 expression profoundly remodels the TME. These analyses revealed a unique unidirectional signaling from CA19-9 producing *Kras* mutant ductal cells to apCAFs to Cd4^+^ T cells. Using a combination of *in vitro* and *in vivo* perturbations, we find that CA19-9 modification of Fbln3 drives Egfr-dependent Il1a and Tgfb elevation in PDAC, promoting apCAF and Treg differentiation which shape the immunosuppressive PDAC microenvironment.

## MATERIALS AND METHODS

### Organoids and monolayer cell line cultures

Detailed procedures for the isolation and propagation of pancreatic ductal organoids cultures have been previously described^34^. Mouse organoid lines, G8529, A5895, and A5897, were derived from Kras^LSL-G12D^; Pdx1-Cre; Rosa26^LSL-rtTA-mKate^; Col1a1^TRE-FUT3-B3GALT5-GFP^ (K;C;R^LSL^;F) mice. Briefly, ducts from pancreata were harvested after enzymatic digestion with 0.012% collagenase XI (Sigma #C7657) and 0.12% dispase (Gibco #17-105-041) in DMEM containing 1% FBS and embedded in 100% growth factor-reduced (GFR) Matrigel (Corning #356231). Mouse organoids were propagated in mouse complete feeding media and maintained at 37°C, 5% CO2. Human patient-derived organoid, hM1F, was obtained from CSHL^35^ and propagated in human complete feeding media and maintained at 37°C in an atmosphere of 5% CO2. Human complete feeding medium was prepared using Advanced DMEM/F12 containing HEPES pH 7.5 (1M, Fisher #50213365), GlutaMAX (1%, Gibco #35050079), B27 supplement (1X, Life Technologies #A3582801), and penicillin/streptomycin (1%, Gibco #15140122), N-acetylcysteine (1.25 mM, Sigma-Aldrich #A9165), nicotinamide (10 mM, Thermo Fischer #125271000), hEGF (50 ng/mL, R&D Systems #236-EG), FGF-10 (100 ng/mL, R&D Systems #345-FG), gastrin (10 nM, R&D #3006), A83-01 (500 nM, Tocris Bioscience #2939), rNoggin (100 ug/mL, Qkine LTD #Qk034), rRSPO1 (10nM Qkine LTD # Qk006), Wnt surrogate (300 pM, ImmunoPrecise Antibodies #N001), and Y-27632 dihydrocholoride (10.5 µM, Tocris Bioscience #1254). Mouse organoid media lacked Wnt3a and Y-27632 and utilized 10% RSPO1 conditioned media and 10% Noggin conditioned media. Mycoplasma testing with the MycoAlert Mycoplasma Detection Kit (Lonza #LT07-318) was performed monthly, and each cell line has been tested at least once after thawing or isolation and retested prior to RNA sequencing and orthotopic transplantation experiments.

Monolayer FC1245 (female) and FC1199 (male) pancreatic cancer cell lines derived from mouse pancreatic tumors of KPC mice on a C57BL/6J pure background and isolated as described previously^33^. Cells were propagated in 10% FBS, 1% P/S in DMEM. Human pancreatic cancer cell line, Capan2, was propagated in 10% FBS, 1% P/S (Gibco #15140122) in RPMI (Gibco #61870036) and maintained at 37°C, 5% CO2.

PanMESO cells were provided by Rolf Brekken^13^. The PanMeso cells were cultured in mesothelial cell media (Medium 199, Gibco #11150059), 10% FBS, 1% penicillin/streptomycin (Gibco #15140122), 3.3 nM mouse epidermal growth factor (Biolegend), 400 nM hydrocortisone (MilliporeSigma), 870 nM zinc-free bovine insulin (MilliporeSigma), and 20 mM HEPES (Fisher #50213365), maintained at 33°C.

### Organoid Conditioned Media

Pancreatic ductal organoids were dissociated into single cells as previously described [44] and seeded as 20 matrigel domes (70ul per dome) in 10cm plates. A total of 13mL of organoid culture medium was added. Conditioned media were collected after 3 days, following 24 hour recovery period post-seeding. To induce CA19-9 expression, organoids were treated with1 ug/ml doxycycline (Sigma #D9891), and conditioned media was collected 3 days after doxycycline treatment. To block CA19-9 expression, organoids were first treated with 10ug/ml CA19-9-blocking antibody HuMab-5B1 (MVT-5873) for 24 hours, followed by treatment with 1ug/ml doxycycline (Sigma #D9891). Conditioned media was then collected 3 days after doxycycline treatment.

### PanMeso Differentiation

PanMeso cells were obtained from Rolf Brekken^13^. 50,000 PanMeso cells were seeded into each well of 6-well plate. After 24 hours, cells were washed three times with 1X PBS and then treated with 3mL of organoid conditioned media for 3 days at 37°C. Total RNA was extracted using TRIzol (Thermo Fisher, #15596018), and quantitative RT-PCR (Table S2) was performed to assess differentiation.

### RNA isolation, cDNA synthesis and Quantitative RT-PCR

RNA was extracted from cell and organoid cultures using TRIzol (Thermo Fisher Scientific #15596018) and RNeasy Mini kit (Qiagen #74104) according to the manufacturer’s instructions. cDNA was synthesized using 1μg of total RNA and SuperScriptTM IV First-Strand Synthesis System (Invitrogen #18091050). The expression of genes was measured by qRT-PCR with Fast SYBR^TM^ Green Master Mix (Applied Biosystems #4385612) on a QuantStudio 6-flex Real time-PCR instrument (Applied Biosystems). For a detailed list of PCR primer sequences, see Table S2. Relative gene expression quantification was performed using the ddCt method with the QuantStudio Real-Time PCR software v1.1 (Applied Biosystems).

### In Vitro Drug and Antibody Treatments

To induce CA19-9 expression, organoids were treated with a final concentration of 1ug/ml doxycycline. For antibody blocking experiments, cells and organoids were treated with 10ug/ml mouse Il1a (BioXcell, #BE0143) or Armenian Hamster IgG control (Leinco, #I-140), 10ug/ml mouse TGFb (BioXcell, #BE0057) or mouse IgG control (Leinco, I-536), 60nM Afatinib or DMSO, 5ug/ml mAb3-5 (Santa Cruz, #sc-33722) or mouse IgG control (Leinco, I-536), fully humanized Fbln3 ab^36^ or human IgG1 control (BioXcell, #BP0297), and 10ug/ml HuMab-5B1 (MVT-5873) or human IgG1 control (BioXcell, #BP0297).

### Lentiviral production

HEK cells were seeded at 6×10^6^ cells per 10cm tissue culture plate. 24 hours later, the cells were 70-80% confluent. Transfection was performed according to manufacturer instructions (Lipofectamine 3000, Invitrogen, #L3000015). Viral supernatant was harvested at 24 hours. One volume of Lenti-X concentrator (Clontech) was added to three volumes of viral supernatant and incubated at 4°C overnight. After centrifugation at 1500xg for 45 minutes at 4°C, the virus pellet was resuspended with appropriate media for transduction.

For infection of monolayer cells, the virus was resuspended in 10% FBS supplemented DMEM containing 10ug/mL polybrene. Selection was initiated 48 hours post-infection with puromycin (2ug/mL, Fisher #HY-B1743A). For infection of organoids, the viral pellet was resuspended in organoid media containing 10ug/ml polybrene and 10.5uM Y-27632 (Tocris Bioscience #1254). Organoid infections were performed following single cell dissociation using spinoculation at 600RCF for 1 hour at room temperature (100,000 cells) followed by resuspension in Matrigel and propagation as organoids in the presence of 10.5uM Y-27632 (Tocris Bioscience #1254) for the first 72 hours. Selection was initiated 48 hours post-infection. Two days after infection, organoids were treated with puromycin (2ug/mL; Fisher #HY-B1743A) for selection.

### Knockdown of *Efemp1* (Fbln3)

To knock down *Efemp1*(Fbln3) expression, DNA sequences encoding shRNAs and a scramble sequence were cloned into pLKO.1 lentiviral vector (Dharmacon, TRCN0000109681). The mouse monolayer cell lines FC1245 and FC1199, organoid lines A5895 and A5897, and human PDO hM1F were transduced with lentivirus as described above.

### Knockout of Fut3

To knockout FUT3, three sgRNAs were co-transduced into Capan-2 cells (RFP-puro) stably expressing Cas9 (Blasticidin). The sgRNA sequences used were: #1:TCCCAGTGGGTCCTCCCGAC; #2:CGATGCCACTGGATCCCCTA; #3:CAGCGGCGCCATGGCCATTG. CA19-9 negative, FUT3 KO cells were sorted using the NS19-9 antibody.

### Animals

Kras^LSL-G12D^;Pdx1-Cre (KC) and Kras^LSL-G12D^;Pdx1-Cre;Rosa26^LSL-rtTA-mKate^;Col1a1^TRE-FUT3-B3GALT5-GFP^ (K;C;R^LSL^;F) mice were generated as previously described^33,37,38^. All animal experiments were conducted in accordance with procedures approved by the IACUC at Salk Institute for Biological Studies. Mice were enrolled into experimental studies between 8 and 14 weeks of age. Mice were administered Dox through drinking water (0.5 - 2mg/ml) supplemented with 10mg/ml Sucrose for study durations up to one week. Dox was administered through chow for experimental durations more than a week, which is designed to deliver 2 - 3mg of Dox per day based on the consumption of 4 - 5g/day per mouse (inotiv, dietTD.01306, Dox 625mg). Therapeutic intervention studies were performed using intraperitoneal injection of 200-1000μg of anti-mouse Il1a (BioXCell, #BE0143) or armenian hamster IgG control (Leinco, #I-140); anti-mouse TGFb (BioXCell, #BE0057) or mouse IgG control (Leinco, I-536); HuMab-5B1 (MVT-5873) or human IgG1 control (BioXCell, #BP0297); anti-mouse Cd8a (Leinco, #C380) or rat IgG control (Leinco, #I-1034); anti-mouse Cd25 (BioXCell, #BE0012) or rat IgG control (Leinco, #R-1379).

### Orthotopic models

FC1245 (female) or FC1199 (male) with or without Fbln3 KD cells were injected orthotopically (100 cells) in 8-12 weeks old C57BL/6J mice (sex matched, n=5/group). Longitudinal tumor volumes were monitored by high resolution ultrasound. Tumors were harvested 5-7 weeks after injection for histologic analyses.

### Immunohistochemistry and immunofluorescence

Tissues were fixed in 10% neutral buffered formalin for 48 hours and embedded in paraffin. Sections were deparaffinized in xylene (2 changes), rehydrated through graded alcohols (2 changes 100% ethanol, 2 changes 95% ethanol, 1 change 80% ethanol, 1 change 70% ethanol) and rinsed in tap water. Antigen retrieval was performed in either 10mM sodium citrate buffer, pH 6.0 or 1mM EDTA, pH8.0 for 20 minutes in a 2100 Retriever (Aptum Biologics), unless otherwise specified. Sections were allowed to cool for 30 minutes and then rinsed in tap water. Slides were assembled in Sequenza slide racks (EMS #71407-1) and washed with 1X TBST for 5 minutes at room temperature. Endogenous peroxidase was quenched with 3% Hydrogen peroxide in TBST for 20 minutes at room temperature. Slides were blocked in 2.5% normal horse serum with 10% normal goat serum for one hour at room temperature and incubated in primary antibody overnight. For antibody dilutions and antigen retrieval conditions, see Table S1. Slides were washed three times in 1X TBST and incubated with horse radish peroxidase-conjugated secondary antibodies for 30 minutes at room temperature. After two additional washes in 1X TBST, slides were disassembled from the cover plates. ImmPACT DAB (Vector #SK-4103-100) was applied for 1 - 5 minutes followed by a water rinse and hematoxylin counterstain for 1 minute. The IHC slides were then dehydrated through graded alcohols (1 change 70% ethanol, 1 change 80% ethanol, 2 changes 95% ethanol, 2 changes 100% ethanol) and 2 changes of xylene. Slides were coverslipped using Omnimount (National diagnostics, #HS-100) and dried overnight in a fume hood. Slides were imaged using Olympus cellSens. The percentage of positive cells were counted in at least five representative high-powered fields from at least three animals per group using QuPath.

### Protein isolation

For monolayer cell lines, tissue culture plates were washed two times with ice cold PBS and lysed on the plate in 1% Triton X-100, 50mM Tris-HCl pH 7.5, 150mM NaCl with 1X protease and phosphatase inhibitor cocktails on ice for 10 minutes. The lysate was transferred to a microcentrifuge tube and clarified at 14,000 RCF for 10 minutes at 4°C, and aliquoted into new tubes for storage at -80°C.

Organoid domes were dislodged using a cell lifter and transferred to a protein LoBind 1.5mL tube (Eppendorf). The organoids were incubated in Cell Recovery Solution (Corning) with 0.5X protease and phosphatase inhibitors on ice for 30 minutes and centrifuged at 800 RCF for 5 minutes at 4°C. The supernatant was discarded. Organoid pellets were snap frozen in liquid nitrogen and stored at -80°C. Organoids were lysed in 1% Triton X-100, 50mM Tris-HCl pH 7.5, 150mM NaCl with 1X protease and phosphatase inhibitor cocktails on ice for 30 minutes, centrifuged at 14,000 RCF for 10 minutes at 4°C, and transferred into new tubes for storage at - 80°C.

### Immunoblot

Protein concentration was measured using BCA assay (Thermo Scientific #23227). Protein lysates were separated on 4-12% Bis-Tris NuPAGE gels (Life technologies) and transferred to PVDF membrane (Millipore). The membranes were probed with pEGFR (Y1068 ; CST #3777), EGFR (CST #71655), p-p65 (NF-kB) (S536; CST #3033), p65 (NF-kB) (CST #8242), pAkt (S473; CST #4060), AKT (CST #4685), pSMAD2 (CST #3108), SMAD2/3 (CST #8685), pERK (T202/Y204; CST #4370), ERK (CST #4695), hFbln3 (RnD #AF6235), CA19-9 (HuMab-5B1 (MVT-5873), and Cofilin (CST #5175) antibodies. For antibody dilutions and additional details, see Table S1. Secondary antibodies used were Peroxidase AffiniPure^TM^ Goat Anti-Mouse IgG (H+L) (Jackson ImmunoResearch #115-035-003), Peroxidase AffiniPure™ Goat Anti-Rabbit IgG (H+L) (Jackson ImmunoResearch #111-035-003), Sheep IgG Horseradish Peroxidase-conjugated Antibody (RnD #HAF016), Goat Anti-Human IgG (ThermoFisher #31410), IRDYE 800CW Goat Anti-Rabbit IgG (Li-Cor #926-32211), and IRDYE 680RD Goat Anti-Mouse IgG (Li-Cor #926-68070) (see Table S1 for antibody dilutions and additional details).

Western blots were developed using either a film processor (Mini-Med 90-AFP Manufacturing Corporation) or Odyssey M (LiCor). Quantification of immunoblots was conducted using either ImageJ or Empiria Studio.

### Single cell dissociation (scRNA-seq)

Pancreata were dissected from KC and KCR^LSL^F mice and finely minced. Pancreata were dissociated in Gentle Collagenase+Hyaluronidase (Stem Cell Technologies #7919) diluted 1:2 to a final volume of 10mL in basal media: Advanced DMEM/F12 (Gibco #12634), 25mM HEPES, Pen/Strep; 1X GlutaMAX (Gibco #35050), 3% BSA; 1mg/mL Soybean Trypsin Inhibitor (Millipore Sigma #93620), and 100ug/mL Dnase (Roche #1128493200)). Samples were incubated on a rotary incubator at 35°C for 20 minutes. Supernatant was collected, centrifuged at 300 RCF for 5 minutes, resuspended in Buffer, and left on ice. 10mL of fresh Gentle Collagenase+Hyaluronidase (Stem Cell Technologies #7919) diluted 1:2 in basal media was added to pancreata and the process was repeated for a total of 4 fractions. For the last fraction, 5U/mL of Dispase (Stem Cell Technologies #7913) in HBSS was used to fully dissociate pancreata. All five fractions were combined, filtered through a 100um cell strainer, centrifuged at 300 RCF for 5 minutes, and resuspended in 5mL of basal media. Viable cell count was performed with Trypan Blue in automated cell counter (Bio-Rad #TC20).

### Single-cell capture, library preparation, and scRNA-seq

Up to 12,000 dissociated cells were loaded per lane on 10X Chromium Next GEM chips. Single-cell capture, barcoding, and library preparation were performed using Chromium Next GEM Single Cell 3’ v3.1 (Dual Index) according to the manufacturer’s instructions (#CG000315). cDNA and libraries were checked for quality on Tapestation (Agilent) and quantified by Qubit Fluorometer (Invitrogen) 297 before sequencing on NextSeq2000 (Illumina).

### Single-cell data processing, quality control, and analysis

The Cell Ranger pipeline (version: cellranger-7.0.0, 10X Genomics) was used to align FASTQs to the mm10 reference (version: refdata-gex-mm10-2020-A, 10X Genomics) for mouse samples and produce digital gene-cell counts matrices. Prior to downstream analysis, ambient RNA contamination was removed using *SoupX* with default parameters. For each sample, the contamination fraction was automatically estimated using the autoEstCont function, and counts matrices were adjusted accordingly. Quality control filtering was performed to retain cells with 50-7,500 detected genes and less than 20% mitochondrial gene content. Each sample was then normalized using log normalization with a scale factor of 10,000, and the 2,000 most variable features were identified using the "vst" method. Doublets were identified and removed using *DoubletFinder*. For each sample, principal component analysis (PCA) was performed using the top 10 principal components. The optimal pK parameter was determined using a parameter sweep without ground-truth classifications, and homotypic doublet proportions were estimated based on preliminary clustering results (resolution 0.8). Doublets were identified assuming an 8.7% doublet formation rate (pN = 0.25, pK = 0.09), and cells classified as doublets were excluded from further analysis. Following doublet removal, samples were integrated using *Harmony* to correct for batch effects. Integrated data were clustered using the Louvain algorithm at a resolution of 0.8 and visualized using UMAP dimensionality reduction. Cluster marker genes were identified using the *FindAllMarkers* function. For differential gene expression analysis between conditions within each cell type, the *FindMarkers* function was used. Manual inspection of acinar-related gene expression was performed to further refine cell type annotations, and cells showing markers of dying cells or residual doublet signatures were removed. Cell-cell communication analysis was performed using *CellChat* with default settings. Gene expression visualization, including violin plots, UMAP plots, and cell type proportion analysis, was performed using the *ShinyCell* package for interactive exploration.

### Single cell dissociation (Flow)

Pancreata were dissected from K;C and K;C;R^LSL^;F mice and finely minced. Pancreata were dissociated at 37°C water bath for 10 minutes in 14mL of Dissociation Buffer: DMEM+GlutaMAX (Gibco #10569), Pen/Strep (Gibco #15140122); 25mM HEPES (Fisher #50213365), 1mg/mL Collagenase XI (Millipore Sigma #C7657), 1mg/mL Soybean Trypsin Inhibitor (Millipore Sigma #93620), and 125ug/mL Dispase; 50ug/mL DNase). Samples were transferred to rotary incubator for 20 minutes at 37C. Supernatant was collected, centrifuged for 5 minutes at 300 RCF and resuspended in Sample Buffer: DMEM+GlutaMAX (Gibco #10569), Pen/Strep, 25mM HEPES, and 10% FBS and left on ice. Fresh Dissociation Buffer was added to sample tubes and process was repeated for a total of 3 fractions. All samples were filtered through a 100um strainer, remaining pieces of tissue were extruded through the strainer with a 5mL syringe plunger. Strainer was washed with Sample Buffer and centrifuged for 5 minutes at 300 RCF. Samples were resuspended in Sample Buffer and viable cell count was performed using Trypan Blue in automated cell counter (Bio-Rad TC20).

Cd4^+^ T cell isolation

Spleens were dissected from OT-II mice and processed through a 70um cell strainer. The cells were then incubated in MACS buffer (1x Dulbecco’s Phosphate Buffered Saline, DPBS; 2mM EDTA, Invitrogen; and 2-5% Normal Rat Serum, Jackson Immunology) with biotinylated antibodies against TER119 (Biolegend #116204), Cd8 (Biolegend #100748), B220 (Biolegend #103204), Cd49b (Biolegend #108904), Cd11b (Biolegend #101204), Cd11c (Biolegend #117304), MHCII (Biolegend #107604), and TCRγδ (BD #553176) (Table S1). Mojosort Streptavidin Nanobeads (Biolegend #480015) were added before placing on an EasySep Magnet (StemCell Technologies #18000) to isolate Cd4^+^ T cells.

### apCAF/Cd4^+^ T cell co-culture assay

KCR^LSL^F mice were treated with Dox. After 3 days, pancreata were dissected and processed through single cell dissociation. Cells were incubated with surface antibodies for 30 minutes at 4°C in 2% FBS in PBS (FACS buffer). After washing, cells were resuspended in FACS buffer with DAPI, Podoplanin (Biolegend #127423), and MHCII (Biolegend #116408). Pdpn^+^MHCII^+^ cells were sorted via BD FACSAria Fusion with a 100um nozzle into HBSS, 50% FBS (Table S1). 20,000 sorted apCAF cells were seeded per well in a 96well U-bottom plate. The next day, cells were incubated with or without 25ug/mL OVA peptide (OVA 323-339) in DMEM, 10% FBS for 4 hours at 37°C. Cells were washed with DPBS 3 times and 50,000 isolated Cd4^+^ T cells were added in DMEM, 10% FBS for 48 hours and collected for flow cytometry.

### Flow Cytometry

Cells were labeled with viability dye (Ghost Dye Red 780, Tonbo; or LIVE/DEAD™ Fixable Violet Dead Cell Stain, Invitrogen #L34955) in PBS for 5 minutes at room temperature prior to antibody staining. Cells were incubated with surface antibodies for 30 minutes at 4°C in 2% FBS in PBS (FACS buffer) (Table S1). Where applicable, a FoxP3 transcription factor staining kit (eBioscience #00-5523-00) was used to fix and permeabilize the cells for 30 minutes at room temperature before incubating with intracellular antibodies for 45 minutes at 4°C. Data acquisition was performed using a LSR II (BD Biosciences), and analysis was performed using FlowJo (TreeStar).

### RNA sequencing and analysis

Organoids in Matrigel were lysed directly with 1 mL of TRIzol reagent 295 (Thermo Fischer #15596018), and total RNA was extracted using RNeasy Mini Kit (Qiagen 296 #74104). RNA quality control was performed on all samples using Qubit Fluorometer (Invitrogen) 297 and TapeStation (Agilent) before RNA-seq analyses. Library construction was conducted using Novaseq 6000 sp reagent kit v 1.5. Samples were sequenced 300 cycles for 150 bp paired end reads.

Paired-end raw FASTQ reads were quality checked with FASTQC (v0.12.1). Poor quality 3’ ends and sequencing adapters were trimmed with Cutadapt (v4.9) and reads shorter than 20nt were removed. Trimmed reads were again checked with FASTQC, then aligned to mouse reference genome (genome mm39 from UCSC and annotations gencode.vM34.basic.annotation.gtf from Gencode) and gene expression quantified with STAR (version 2.7.11b).

Second strand expression was extracted from STAR quantification for all samples to generate a read matrix. DESeq2 was used to normalize read counts and find differentially expressed genes between groups, which is defined as padj < 0.05 and |log2FC| > 1.

### Statistics

Statistical tests were performed using GraphPad Prism 11 software and are reported for each experiment in their corresponding figure legends. All graphs plot individual data points to demonstrate the n values for each condition with the bar representing the mean and the error bars representing the standard deviation. p values are reported as * p<0.05, ** p<0.01, *** p<0.001, **** p<0.0001.

## RESULTS

### CA19-9 induces TME remodeling

Previous work examining PDAC TME remodeling has often been performed in GEMMs harboring *Kras* and *p53* mutations, which recapitulate hallmark features of human disease. However, these studies have been evaluated in a CA19-9 negative context due to lack of endogenous production of this glycan in mice, despite CA19-9 representing a clinically central biomarker and mediator of PDAC pathogenesis. To dissect the contributions of CA19-9 to TME remodeling, we utilized ***K****ras*^LSL-G12D^;*Pdx1-**C**re*;***R****osa26***^L^**^SL-rtTA-mKate^;*Col1a1*^TRE-**F**UT3-B3GALT5-GFP^ (KCR^LSL^F) mice^33^. Prior Dox-inducible CA19-9 modeling in the setting of wildtype *Kras* demonstrated that CA19-9 expression is sufficient to induce pancreatitis, a major PDAC risk factor. Together with *Kras* mutation, 2-4 weeks of CA19-9 induction led to dramatic expansion of pancreatic intraepithelial neoplasia (PanIN) and early emergence of invasive carcinoma, both of which were accompanied by substantial remodeling of the stromal and immune TME^33^. We find that this TME remodeling is rapid, occurring within 72 hours (**Fig. 1A, S1A**) and is characterized by loss of normal acinar architecture (**Fig. 1B**), early- and late-stage PanIN lesions (**Fig. 1B**), collagen deposition (**Fig. 1C**), epithelial and TME proliferation (**Fig. 1D**), and expansion of CAFs (Pdpn^+^) (**Fig. 1E**). This TME remodeling persisted for the lifespan of the mice (**Fig. S1A**). TME remodeling was not induced by an anti-CA19-9 immune response as depletion of Cd8^+^ T cells did not impact CA19-9 induced TME remodeling (**Fig. S1B-C**), which is also consistent with a lack of CA19-9 autoantibodies and absence of non-pancreatic phenotypes previously observed from our previous work^33^. Together, these data reveal that the TME remodeling is not induced by autoimmunity against CA19-9 and instead represents a previously unrecognized contribution of aberrant glycosylation to the immunosuppressive features of the PDAC TME.

**Figure 1.**
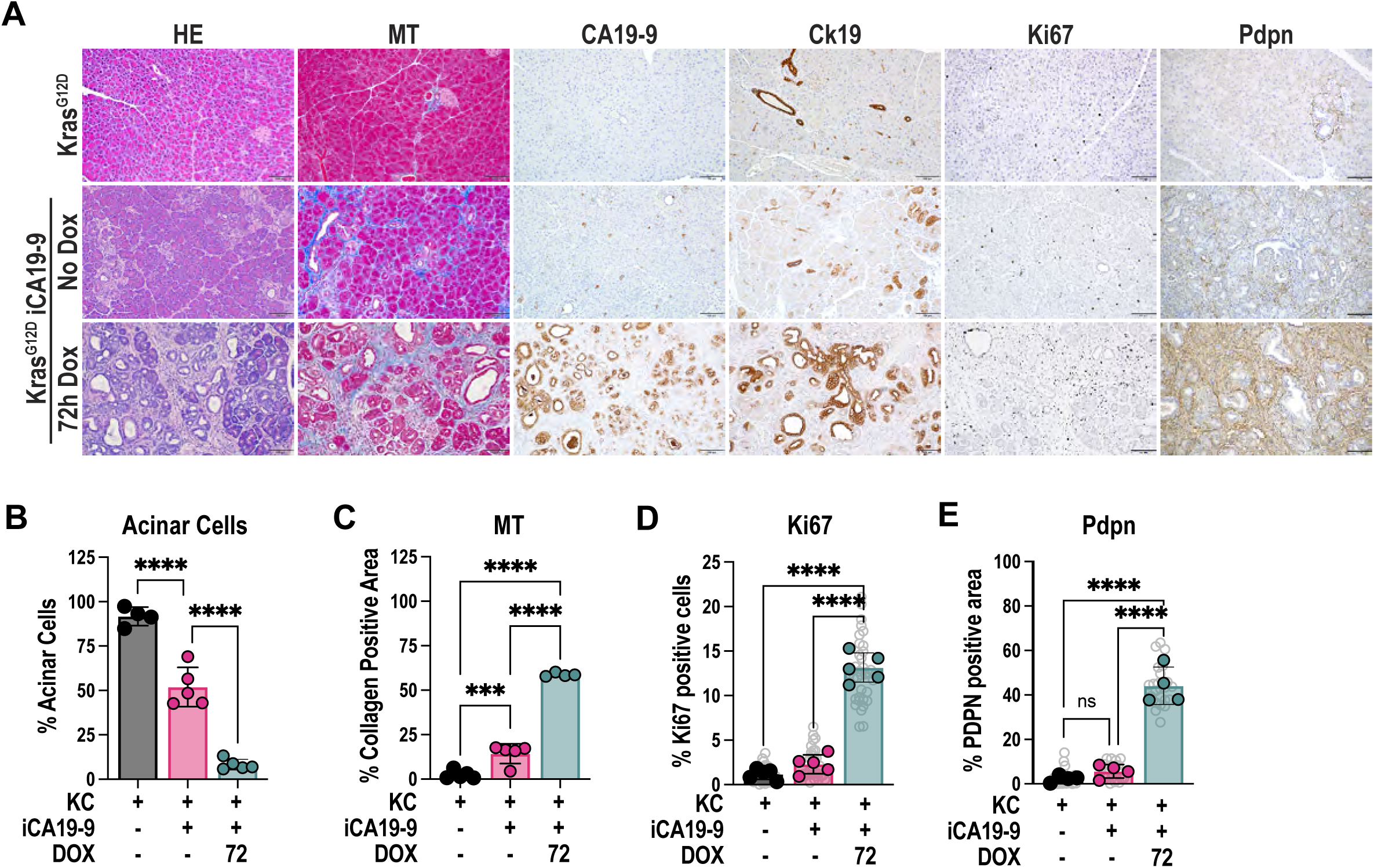
**A)** Histologic evaluation of KC and KCR^LSL^F pancreata after the treatment with Dox by hematoxylin and eosin (HE) staining and Masson’s trichrome staining (MT) (blue=collagen deposition). CA19-9, cytokeratin19 (Ck19), Ki67, and podoplanin (Pdpn) expression evaluated by IHC. Scale bars = 100 um **B)** Acinar cell histological quantification as a percentage of total tissue area in KC and KCR^LSL^F pancreata after the treatment with Dox. Bar graphs represent the mean, and error bars represent the standard deviation. n=5. Each data point represents a measurement from an individual mouse. Statistical analysis by ordinary one-way ANOVA. **C)** Collagen area (blue) histological quantification as a percentage of total tissue area in KC and KCR^LSL^F pancreata after treatment with Dox. Bar graphs represent the mean, and error bars represent the standard deviation. n=5. Each data point represents a measurement from an individual mouse. Statistical analysis by ordinary one-way ANOVA. **D)** Ki67 positivity via histological quantification as a percentage of total cells in single FOVs in KC and KCR^LSL^F+Dox pancreta. Bar graphs represent the mean, and error bars represent the standard deviation. n=5. Each grey circle represents one FOV. Each solid data point represents the average of all FOVs/mouse. Statistical analysis by ordinary one-way ANOVA. **E)** Pdpn area histological quantification as a percentage of total tissue area in single FOVs in KC and KCR^LSL^F pancreata after treatment with Dox. Bar graphs represent the mean, and error bars represent the standard deviation. n=4. Each grey circle represents one FOV. Each solid data point represents the average of all FOVs/mouse. Statistical analysis by ordinary one-way ANOVA.

### apCAFs and Tregs expand in CA19-9 positive Kras-mutant pancreata

To map the CA19-9 induced changes to the TME, we performed single-cell RNA sequencing (scRNA-seq) on pancreata from CA19-9^pos^ Dox-treated (3 days) and untreated CA19-9^low^ KCR^LSL^F mice as well as CA19-9^neg^ ***K****ras*^LSL-G12D^;*Pdx1-**C**re* (KC) controls (**Fig. 2A, S2A**). No sex-based differences were observed and all cell types identified were distributed across the biologic replicates of each group (**Fig. S2A**). As anticipated, KC pancreata were predominantly composed of acinar cells (**Fig. 2A-B**). Untreated KCR^LSL^F pancreata, as previously reported, exhibit a small amount of CA19-9 leakage, resulting in an intermediate phenotype with partial TME remodeling, though with reduced representation of stromal and immune populations relative to their Dox-treated counterparts (**Fig. S2A**). In contrast, CA19-9^pos^ KCR^LSL^F pancreata exhibited marked expansion of ductal, CAF, and immune compartments (**Fig. 2B, S2B-C**).

**Figure 2.**
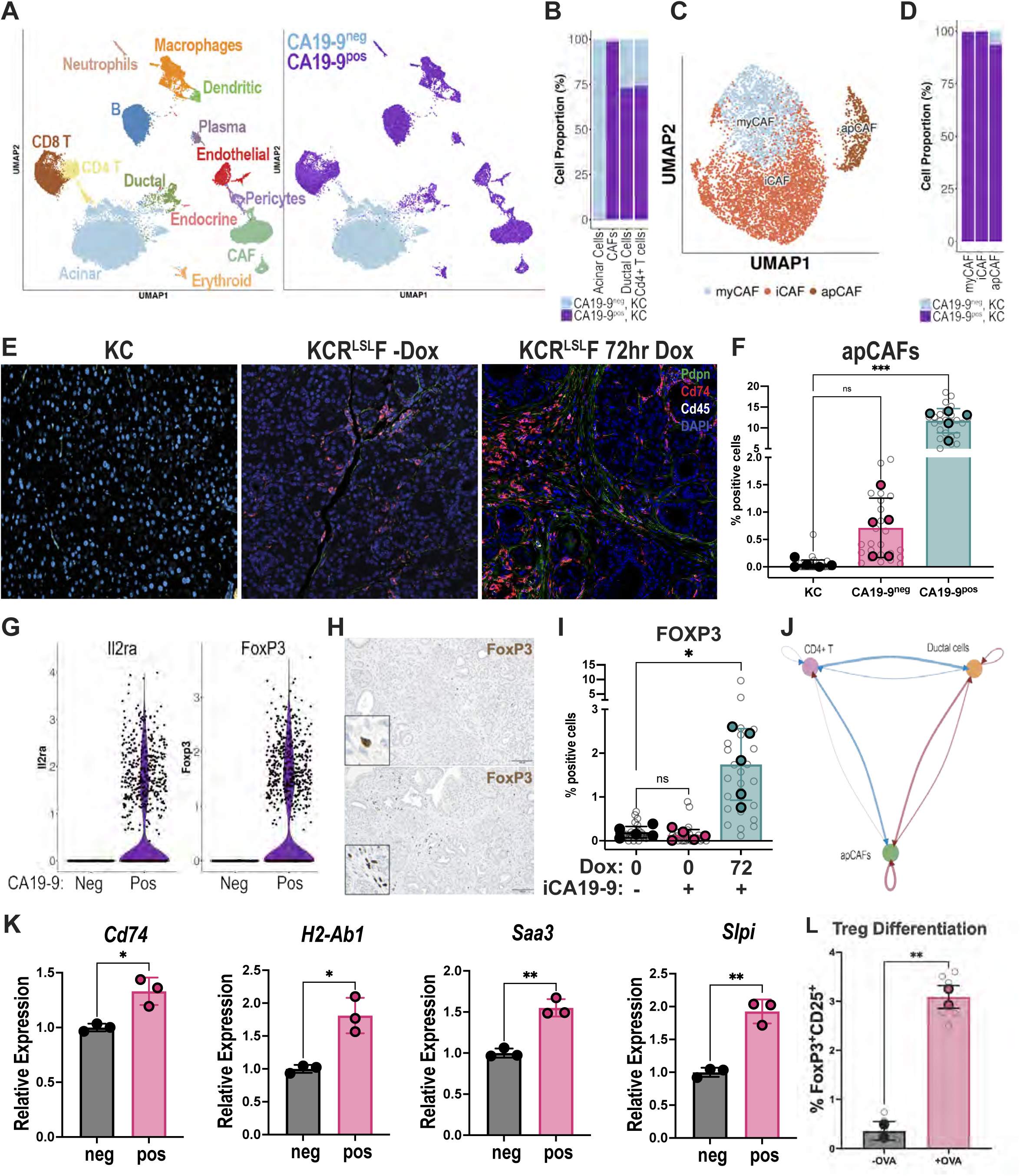
**A)** Uniform manifold approximation and projection (UMAP) visualization of pooled 12,000 single cells per mouse pancreata from KC and KCR^LSL^F treated with Dox (n=3, 5 respectively). Cells are colored by cell type as annotated by literature-defined gene signatures. **B)** Cell proportion graphs denoting relative percentage of Acinar, CAFs, Ductal, and Cd4+ T cells in CA19-9^pos^ KC vs CA19-9^neg^ KC cells. **C)** UMAP projection of CAF subtypes of pooled 12,000 single cells per pancreata from KC and KCR^LSL^F+Dox (n=3, 5 respectively). **D)** Cell proportion graph from scRNAseq denoting relative percentage of myofibroblastic CAF (myCAF), inflammatory CAF (iCAF) and antigen-presenting CAF (apCAF) in CA19-9^pos^ KC vs CA19-9^neg^ KC cells. **E)** Immunofluorescent histochemistry evaluation of apCAFs (Pdpn^+^Cd74^+^Cd45^-^) in KC and KCR^LSL^F mice treated with Dox. Scale bar = 23um **F)** Quantification of apCAFs as a percentage of total cells in single FOVs in KC and KCR^LSL^F mice treated with Dox. Bar graphs represent the mean, and error bars represent the standard deviation. n=5. Each grey circle represents one FOV. Each solid data point represents the average of all FOVs/mouse. Statistical analysis by Kruskal-Wallis test. **G)** Violin plots of Il2ra (Cd25) and FoxP3 gene expression of single cells in KC and KCR^LSL^F+Dox pancreata. **H)** Histologic evaluation of KC and KCR^LSL^F pancreata by FoxP3 IHC staining. Scale bars = 100um **I)** FoxP3 histological quantification as a percentage of total cells in single FOVs in KC and KCR^LSL^F pancreata. Bar graphs represent the mean, and error bars represent the standard deviation. n=5. Each grey circle represents one FOV. Each solid data point represents the average of all FOVs/mouse. Statistical analysis by Kruskal-Wallis test. **J)** CellChat projection cell-cell signaling among Ductal, apCAF, and Cd4^+^ T cells. Red lines indicate upregulation of cell signaling and blue lines indicated downregulation comparing CA19-9^pos^ and CA19-9^neg^ conditions. Lines on the exterior of the graph denote out-bound signaling in a clockwise fashion and interior lines are counter-clockwise. **K)** PanMESO cells were treated with conditioned media (CM) from K;C;R^LSL^;F organoids with or without prior doxycycline (Dox) treatment for 72hrs. Cells were harvested and analyzed by qPCR for apCAF markers: Cd74, H2-Ab1 (MHCII), Saa3, and Slpi. *n=3*. Data shown as mean ± SD. Statistical analysis by unpaired, two-tailed Welch’s corrected t test. **L)** Flow cytometry quantification of Treg (Cd25+FoxP3+) differentiation of OT-II Naive Cd4^+^ T cells when co-cultured with KCR^LSL^F sorted apCAFs for 48 hours in the presence of OVA antigen. Open circles represent mean of 3 technical replicates per mouse and bar graphs represent mean of biological replicates, and error bars represent the standard deviation. Statistical analysis by unpaired, two-tailed Welch’s corrected t test.

Within the CAF compartment, we identified myCAF (*Acta2*^+^*Inhba*^+^*Tagln*^+^), iCAF (*Ly6c1*^+^*Cd34*^+^*Cxcl12*^+^), and apCAF (*Cd74*^+^*H2-Ab1*^+^*Slpi*^+^) subtypes, consistent with prior PDAC studies^11,19,39^ (**Fig. 2C-D, S2D, S3B**). Immunohistochemical and flow cytometric analysis confirmed significant enrichment of apCAFs (Pdpn^+^Cd74^+^Cd45^-^) in CA19-9^pos^ pancreata compared to both untreated KCR^lsl^F and KC controls (**Fig. 2E-F, S3B**). Prior studies observed that apCAFs express MHC class II (*H2-ab1*) and present antigens to Cd4^+^ T cells, thereby promoting antigen-dependent Treg induction, contributing to immunosuppression^11,13^. Consistent with this, Treg-associated transcripts Il2ra (Cd25) and Foxp3 were elevated in the Cd4 T cell cluster of CA19-9^pos^ pancreata relative to KC controls, while markers of other Cd4^+^ T cell subtypes were not highly represented (**Fig. 2G, S2E**). We histologically confirmed an increase in Foxp3 positive cells after 72 hours of CA19-9 induction (**Fig. 2I**). To investigate this paracrine signaling network further, we utilized CellChat^40^, which computationally infers cell-to-cell interactions from scRNA-seq data. We found bi-directional signaling between ductal cells and apCAFs and unidirectional signaling from apCAFs to Cd4^+^ T cells strengthened in response to CA19-9 elevation (**Fig. 2J, S2F-G**). To mechanistically dissect this nexus, we first interrogated CA19-9 driven apCAF differentiation. We treated pancreatic mesothelial (PanMeso) cells^13^ with conditioned media (CM) from KCR^LSL^F mouse organoids with or without Dox treatment or non-conditioned media with Dox. CA19-9^pos^ media significantly upregulated MHC class II and apCAF gene expression (H2-Ab1, Cd74, Saa3, Slpi) (**Fig. 2K**). To assess apCAF-induced Treg differentiation *in vitro*, we sorted apCAFs (Pdpn^⁺^Cd74^⁺^Cd45⁻) from CA19-9^pos^ KCR^LSL^F mouse pancreata following 3 days of Dox treatment and co-cultured them with naïve Cd4^+^ T cells from OT-II mouse spleens for 48 hours after pre-loading OVA peptide. Treg differentiation (Cd25^+^FoxP3^+^) in co-culture was induced in an antigen-dependent manner and was not observed when OT-II Cd4^+^ T cells alone were treated with CA19-9^neg^ or CA19-9^pos^ KCR^LSL^F organoid CM (**Fig. 2L, S3C)**. Together, these analyses reveal that CA19-9 expression in *Kras*-mutant CA19-9^pos^ pancreata rapidly reconstitutes cellular ecosystems characteristic of PDAC, including specialized fibroblast states and regulatory immune circuits, thereby mirroring early human pancreatic tumorigenesis and immunosuppressive conditioning.

### CA19-9-dependent changes to apCAFs and Tregs

A major benefit of the TetOn system to activate CA19-9 production is the ability to also perform pulse/chase analyses. However, despite cessation of Dox administration, eGFP (downstream of Tre promoter) and CA19-9 expression (from FUT3 and B3GALT5 expression downstream of Tre promoter) was maintained even after 24 weeks (**Fig. S3D**), despite the use of reverse osmosis water. These data suggest that the reversible nature of the TetOn system is not functional in the pancreas despite demonstration of reversibility in other organs (e.g., liver)^41^. Therefore, to investigate the direct role of CA19-9 on TME remodeling we utilized CA19-9 antibody blockade (5B1) for *in vivo* intervention studies. In human patients with PDAC enrolled in clinical trials using this same antibody clone^42^, efficacy of the antibody treatment as a mono therapy or in combination with chemotherapy was limited to patients with less than 2,500U/mL of CA19-9 in their serum due to target-mediated degradation in circulation. Consistent with this, efficacy therapeutic antibody blockade in these mice was limited when initiated after CA19-9 induction (**Fig. S4A**). These data highlight the challenge of effective blockade of this high abundance, circulating target and the necessity of dissecting effector pathways of this paracrine signaling axis to identify additional nodes for interception. Pre-treatment of KCR^LSL^F mice with CA19-9 blocking antibody attenuated TME remodeling and disease progression following subsequent treatment with Dox for 72 hours (**Fig 3A**), maintaining the acinar cell population and normal cellular architecture as well as limiting collagen deposition, acinar to ductal metaplasia (Ck19^+^), proliferation (Ki67^+^), and fibroblast expansion (Pdpn^+^) (**Fig. 3B-E**). We also observe reduced apCAF and Treg differentiation (**Fig. 3F-G, S4B-C**), suggesting that the TME remodeling is directly mediated by CA19-9. To test the impact of CA19-9 elevation in the ductal compartment on apCAF differentiation *in vitro*, we treated KCR^LSL^F organoids with Dox in the presence of CA19-9 blocking antibody or isotype control. Following treatment of panMeso cells with CM, we found differentiation to apCAFs was significantly decreased in the presence of CA19-9 blocking antibody (**Fig. 3H**). Together, these findings demonstrate that CA19-9 antibody pre-treatment protects normal pancreatic histology, mitigates stromal remodeling, and reduces apCAF and Treg differentiation, suggesting a direct role of this glycan in TME remodeling.

**Figure 3.**
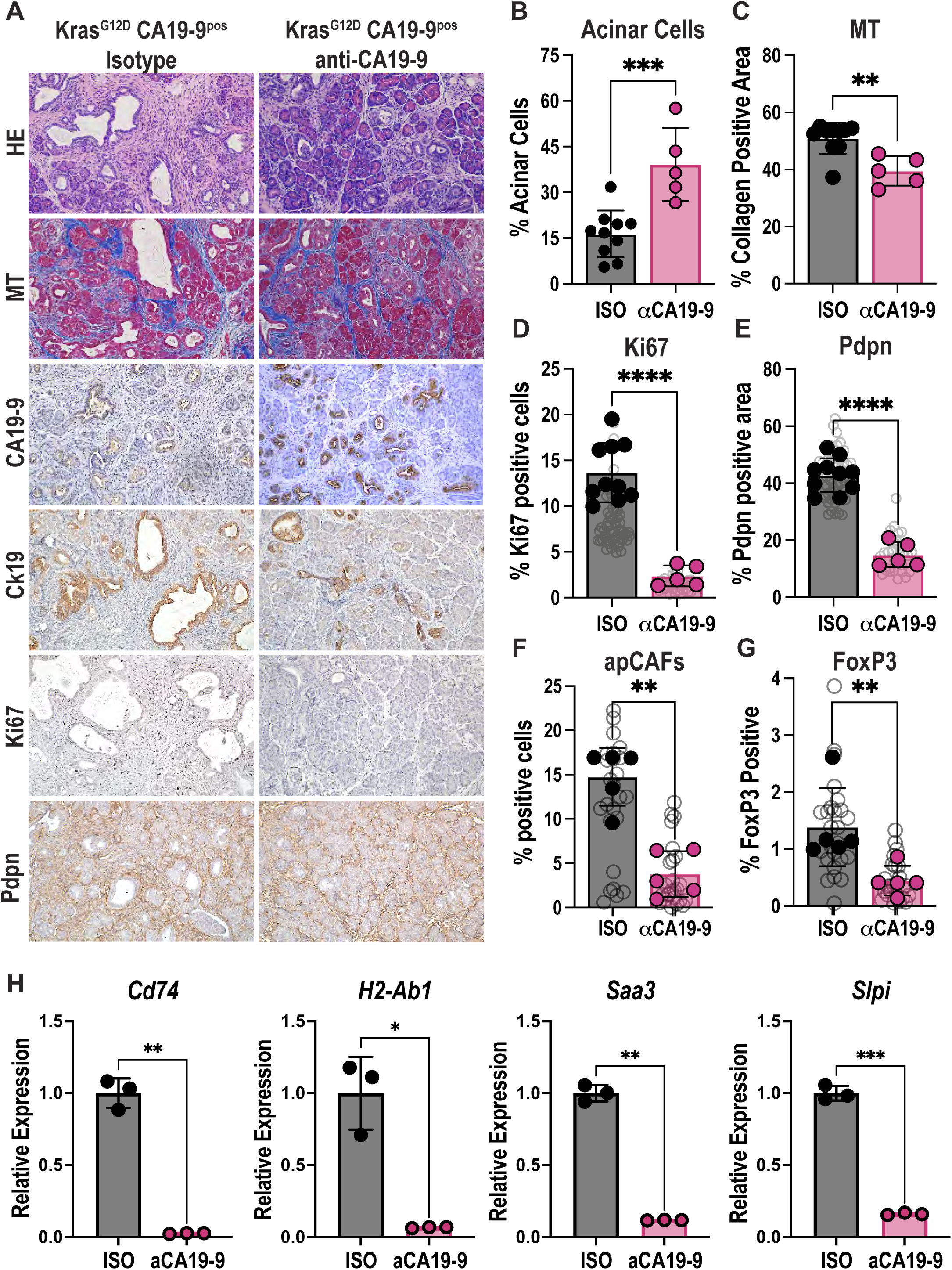
**A)** Histologic evaluation of KCR^LSL^F pancreata after treatment of Dox and anti-CA19-9 antibody blockade compared to isotype control by HE and MT staining. CA19-9, CK19, Ki67, and PDPN expression evaluated by IHC. Scale bars = 100um. **B)** Acinar cell histological quantification as a percentage of total tissue area in KCR^LSL^F pancreata after the treatment with Dox with anti-CA19-9 antibody blockade compared to isotype control (n=5, 10 respectively). Bar graphs represent the mean, and error bars represent the standard deviation. Each data point represents a measurement from an individual mouse. Statistical analysis by unpaired t test. **C)** Collagen area histological quantification as a percentage of total tissue area in KCR^LSL^F pancreata after the treatment with Dox with anti-CA19-9 antibody blockade compared to isotype control (n=5, 10 respectively). Bar graphs represent the mean, and error bars represent the standard deviation. Each data point represents a measurement from an individual mouse. Statistical analysis by unpaired, two-tailed Welch’s corrected t test. **D)** Ki67 positivity via histological quantification as a percentage of total cells in single FOVs in KCR^LSL^F pancreata after the treatment with Dox with anti-CA19-9 antibody blockade compared to isotype control (n=5, 10 respectively). Bar graphs represent the mean, and error bars represent the standard deviation. n=5. Each grey circle represents one FOV. Each solid data point represents the average of all FOVs/mouse. Statistical analysis by unpaired, two-tailed Welch’s corrected t test. **E)** PDPN area histological quantification as a percentage of total tissue area in single FOVs in KCR^LSL^F pancreata after the treatment with Dox with anti-CA19-9 antibody blockade compared to isotype control (n=5, 10 respectively). Bar graphs represent the mean, and error bars represent the standard deviation. Each grey circle represents one FOV. Each solid data point represents the average of all FOVs/mouse. Statistical analysis by unpaired, two-tailed Welch’s corrected t test. **F)** Quantification of apCAFs as a percentage of total cells in single FOVs in KCR^LSL^F pancreata after the treatment with Dox with anti-CA19-9 antibody blockade compared to isotype control (n=5). Bar graphs represent the mean, and error bars represent the standard deviation. Each grey circle represents one FOV. Each solid data point represents the average of all FOVs/mouse. Statistical analysis by unpaired, two-tailed Welch’s corrected t test. **G)** FoxP3 positivity via histological quantification as a percentage of total cells in single FOVs in KCR^LSL^F pancreata after the treatment with Dox with anti-CA19-9 antibody blockade compared to isotype control (n=5). Bar graphs represent the mean, and error bars represent the standard deviation. Each grey circle represents one FOV. Each solid data point represents the average of all FOVs/mouse. Statistical analysis by unpaired, two-tailed Welch’s corrected t test. **H)** PanMESO cells were treated with CM from K;C;R^LSL^;F organoids pre-treated with both Dox and CA19-9-blocking antibody or an IgG isotype control (ISO) for 72 hours. Cells were harvested and analyzed by qPCR for apCAF markers: Cd74, H2-Ab1 (MHCII), Saa3, and Slpi. *n=3*. Data shown as mean ± SD. Statistical analysis by unpaired, two-tailed Welch’s corrected t test.

### IL1a and TGFb expression is induced by CA19-9 and blockade prevents *in vitro* apCAF differentiation

Il1a and Tgfb have been implicated as essential cues for apCAF induction^13^. To determine how CA19-9 elevation contributes to apCAF induction, we investigated changes to signaling networks in KCR^LSL^F organoids using RNA-seq. Gene set enrichment analyses revealed that CA19-9 enhanced Nfkb and Tgfb signaling pathways (**Fig. 4A**). As expected, based on our previous findings, we also find increased enrichment of Egfr signaling following CA19-9 elevation (**Fig. 4A**). The transcript levels of *Il1a* and *Tgfb* were also significantly upregulated followed by CA19-9 elevation (**Fig. 4B, Fig. S5A**), which we validated by qPCR and at the protein level in organoid conditioned media (**Fig. 4C**). Il1a and Tgfb proteins were similarly upregulated *in vivo* in lysates from CA19-9^pos^ KCR^LSL^F pancreata (**Fig. 4D**). To test CA19-9 mediated induction of IL1A and TGFB in a human setting, we used CRISPR/Cas9-mediated knockout of *FUT3* to generate CA19-9^low/neg^ Capan2 cells, which was confirmed by immunoblotting and ALPHA assays (**Fig. S5B-C**). We found that CA19-9^pos^ Capan2 cells with control sgRNAs exhibited higher *IL1A* and *TGFB* transcript levels than CA19-9^low/neg^ cells (**Fig. S5D**). To test the impact of Il1a and Tgfb loss on CA19-9-induced apCAF differentiation, PanMeso cells were treated with KCR^LSL^F organoid CM following Dox induction with or without anti-Il1a and anti-Tgfb antibody treatment. Recombinant Il1a and Tgfb was used as a positive control (**Fig. S5E**). Combined depletion of Il1a and Tgfb from CA19-9^pos^ KCR^LSL^F organoid CM decreased PanMeso apCAF differentiation (**Fig. 4E**). These data show that CA19-9 elevation increases Il1a and Tgfb levels in both mouse and human cell line and organoid models as well as in mouse pancreatic lysates. Further, we demonstrate that apCAF differentiation is Il1a and Tgfb dependent *in vitro*.

**Figure 4.**
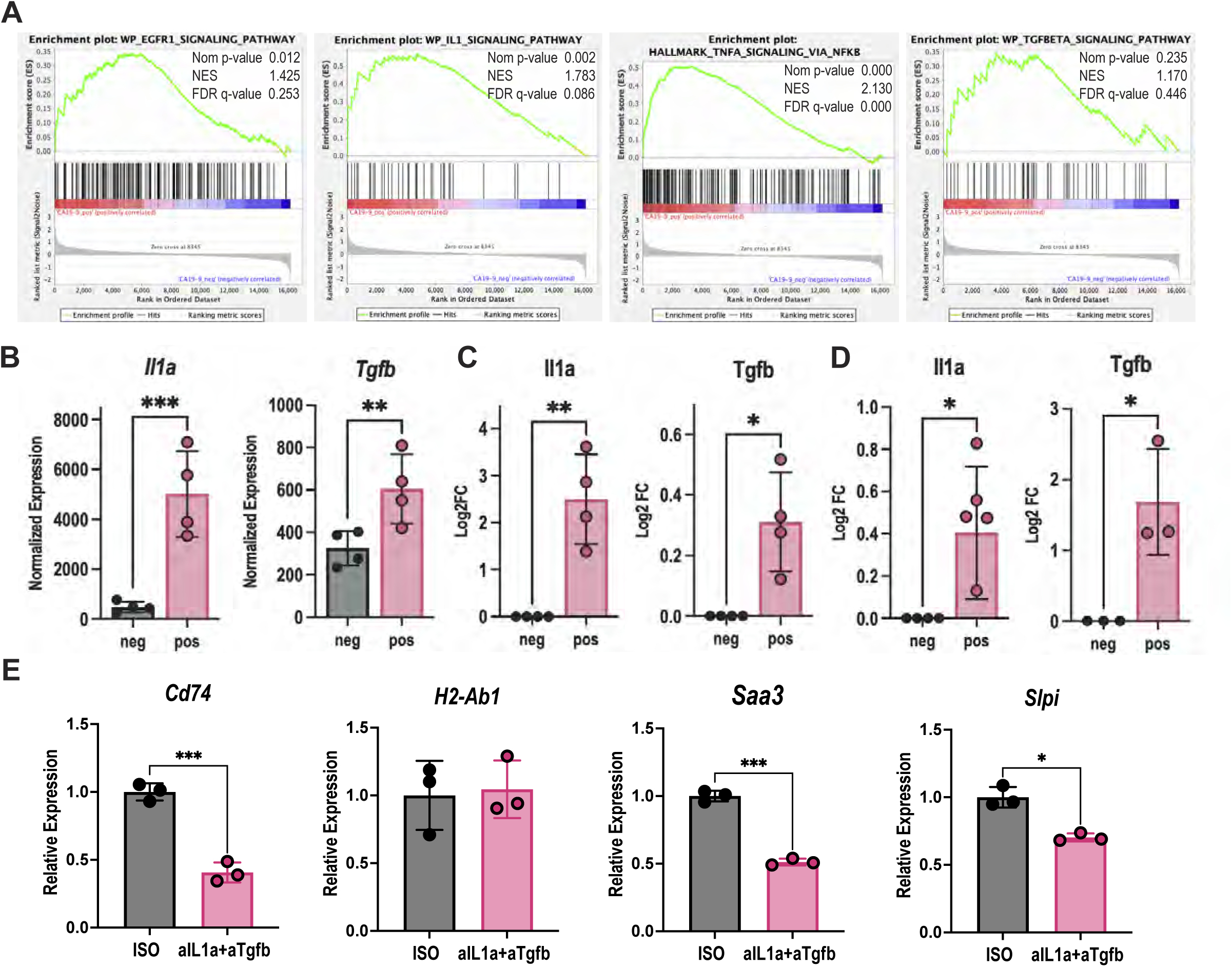
**A)** Gene Set Enrichment Analysis (GSEA) of K;C;R^LSL^;F organoids showing enrichment in EGFR1, IL1A, TNFA VIA NFKB, and TGFB signaling gene sets. NES, normalized enrichment score; FDR: false discovery rate. **B)** RNA-seq analysis of Il1a and Tgfb in KCR^LSL^F CA19-9^pos^ and CA19-9^neg^ organoids (24 hours Dox treatment, 100ng/ml). Statistical analysis by paired, log normal t test. **C)** ELISA protein quantification of IL1a and TGFb in conditioned media from KCR^LSL^F CA19-9^pos^ and CA19-9^neg^ organoids. Statistical analysis by paired t test. **D)** ELISA protein quantification of IL1a and TGFb in tissue lysate from KCR^LSL^F CA19-9^pos^ and CA19-9^neg^ pancreata. Statistical analysis by unpaired t test. **E)** PanMESO cells were treated with CM from K;C;R^LSL^;F organoids pre-treated with both Dox and neutralizing antibodies against IL1A and TGFB or IgG isotype controls (ISO) for 72 hours. Cells were harvested and analyzed by qPCR for apCAF markers: Cd74, H2-Ab1 (MHCII), Saa3, and Slpi. *n=3*. Data shown as mean ± SD. Statistical analysis by unpaired, two-tailed Welch’s corrected t test.

### Blockade of Il1a and Tgfb in mice reduces apCAF and Treg differentiation

To investigate the contribution of Il1a and Tgfb to CA19-9-induced TME remodeling i*n vivo*, we pre-loaded KCR^LSL^F mice with blocking antibodies prior to 3 days of Dox treatment. Blockade of Il1a or Tgfb alone partially attenuated stromal expansion, whereas combined Il1a and Tgfb Ab treatment significantly and substantially protected acinar cell abundance (**Fig. 5A, S6A**) and reduced collagen deposition (**Fig. 5B, S6B**). Cell proliferation (Ki67^+^) decreased significantly with anti-Il1a monotherapy and combination blockade, whereas anti-Tgfb monotherapy had no effect compared to isotype control (**Fig. 5C, S6C**). CAFs (Pdpn^+^), a principal stromal constituent in PDAC, were modestly reduced by single-agent antibody treatment but were substantially depleted under combinatorial targeting (**Fig. 5D, S6D**). apCAF differentiation (Pdpn^+^Cd74^+^Cd45^-^), known to be driven by Il1a and Tgfb, was reduced by both single-agent treatments and significantly diminished upon combined blockade (**Fig. 5E, S6E**). As expected, given that tumor-derived Il1a promotes Treg chemotaxis^43^ and Tgfb regulates Treg differentiation and expansion^44^, we observed significant reductions of FoxP3⁺ Tregs upon both monotherapy and combined Il1a and Tgfb Ab treatment arms (**Fig. 5F, S6F**). Collectively, these findings suggest that CA19-9-mediated upregulation of Il1a and Tgfb drives apCAF and Treg differentiation in mice.

**Figure 5.**
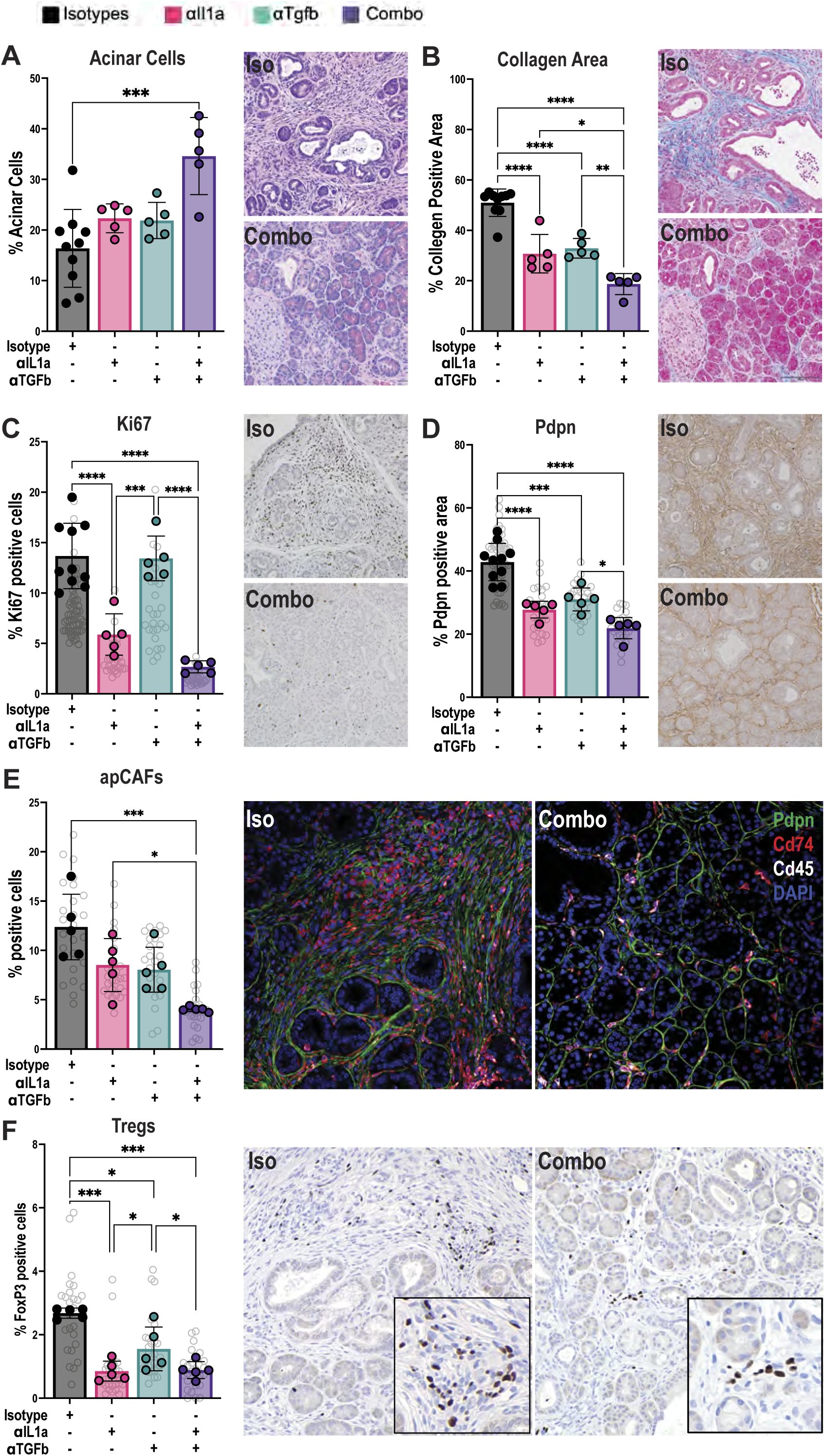
**A)** Representative H&E of combination and isotype arms and acinar cell quantification of all arms as a percentage of total tissue area in KCR^LSL^F pancreata after the treatment with Dox with anti-Il1a, -Tgfb antibody blockade, individually and in combination, compared to isotype control (n=5, 5, 5, 10 respectively). Bar graphs represent the mean, and error bars represent the standard deviation. Each data point represents a measurement from an individual mouse. Statistical analysis by ordinary one-way ANOVA. **B)** Representative MT of combination and isotype arms and collagen area quantification of all arms as a percentage of total tissue area in KCR^LSL^F pancreata after the treatment with Dox with anti-Il1a, -Tgfb antibody blockade, individually and in combination, compared to isotype control (n=5, 5, 5, 10 respectively). Bar graphs represent the mean, and error bars represent the standard deviation. Each data point represents a measurement from an individual mouse. Statistical analysis by ordinary one-way ANOVA. **C)** Representative Ki67 IHC of combination and isotype arms and quantification of all arms as a percentage of total cells in single FOVs in KCR^LSL^F pancreata after the treatment with Dox with anti-Il1a, -Tgfb antibody blockade, individually and in combination, compared to isotype control (n=5, 5, 5, 10 respectively). Bar graphs represent the mean, and error bars represent the standard deviation. n=5. Each grey circle represents one FOV. Each solid data point represents the average of all FOVs/mouse. Statistical analysis by ordinary one-way ANOVA. **D)** Representative Pdpn IHC of combination and isotype arms and quantification of all arms as a percentage of total tissue area in single FOVs in KCR^LSL^F pancreata after the treatment with Dox with anti-Il1a, -Tgfb antibody blockade, individually and in combination, compared to isotype control (n=5, 5, 5, 10 respectively). Bar graphs represent the mean, and error bars represent the standard deviation. n=5. Each grey circle represents one FOV. Each solid data point represents the average of all FOVs/mouse. Statistical analysis by ordinary one-way ANOVA. **E)** Representative apCAF IF of combination and isotype arms and quantification of all arms as a percentage of total cells in single FOVs in KCR^LSL^F pancreata after the treatment with Dox with anti-Il1a, -Tgfb antibody blockade, individually and in combination, compared to isotype control (n=5, 5, 5, 10 respectively). Bar graphs represent the mean, and error bars represent the standard deviation. n=5. Each grey circle represents one FOV. Each solid data point represents the average of all FOVs/mouse. Statistical analysis by ordinary one-way ANOVA. **F)** Representative FoxP3 IHC of combination and isotype arms and quantification of all arms as a percentage of total cells in single FOVs in KCR^LSL^F pancreata after the treatment with Dox with anti-IL1a, -TGFb antibody blockade, individually and in combination, compared to isotype control (n=5, 5, 5, 10 respectively). Bar graphs represent the mean, and error bars represent the standard deviation. n=5. Each grey circle represents one FOV. Each solid data point represents the average of all FOVs/mouse. Statistical analysis by ordinary one-way ANOVA.

### Fbln3-dependent Il1a and Tgfb production mediates apCAF differentiation

We previously showed that CA19-9 modification of Fbln3 drives hyperactivation of the Egfr pathway in pancreatitis^33^. EGFR signaling is frequently upregulated and aberrantly activated in PDA, particularly during acinar-to-ductal metaplasia, where it drives tumor initiation and progression^45^. Immunohistochemical analysis of human pancreatic tissue revealed that FBLN3 is enriched in ductal epithelial cells and is significantly upregulated in PDAC compared to normal pancreata (**Fig. S7A**). To further assess clinical relevance, we analyzed NanoString GeoMx spatial transcriptomic profiles from treatment-naïve primary human PDAC specimens^46^ and found that *FBLN3* expression is significantly elevated in advanced-stage PDAC (**Fig. S7B**).To identify signaling pathways directly regulated by Fbln3, we knocked down *Fbln3* in KCR^LSL^F organoids (**Fig. S7C**) and performed RNA-seq. Transcriptomic and pathway analysis revealed that loss of *Fbln3* resulted in significant suppression of an Egfr signaling gene set in a CA19-9 dependent manner, consistent with our prior findings (**Fig. S7D-E**). To determine which cell populations can respond to Fbln3-mediated Egfr signaling, we analyzed scRNA-seq data and found that Egfr expression was enriched in ductal epithelial cells and CAFs (**Fig. S7F**). Notably, apCAFs expressed little to no Egfr compared with iCAFs and myCAFs (**Fig. S7G**). Analysis of human scRNA-seq data revealed that EGFR expression was not detected in apCAFs (**Fig. S7G)**. Altogether, these data suggest that Fbln3 does not directly act on apCAFs through Egfr but instead regulates their differentiation indirectly through induction of other epithelial-derived factors. Given that Egfr pathway activation can drive expression of Il1a and Tgfb in other cell types^47^, we asked if Fbln3 contributes to CA19-9-mediated Il1a and Tgfb production and subsequent TME remodeling.

Given prior studies demonstrating that epithelial-derived Il1a and Tgfb drives apCAF differentiation^13^, we examined whether Fbln3 regulates the expression of these cytokines. RNA-sequencing analysis revealed that *Fbln3* knockdown (KD) significantly suppressed Nfkb, Tgfb, and Il1a signaling pathways (**Fig. 6A, S7E**). We confirmed that levels of p-p65 (Nfkb) and pSmad2 along with p-Egfr levels were decreased following *Fbln3* KD (**Fig. 6B, S7H**). Correspondingly, Il1a and Tgfb gene and protein expression levels were significantly decreased in *Fbln3* KD organoids (**Fig. 6D-E, S7I**). Antibody-blockade of Fbln3 also decreased Il1a and Tgfb expression as well as levels of pEgfr, pStat3, p-p65, and pSmad2 (**Fig. S7J-K**).

**Figure 6.**
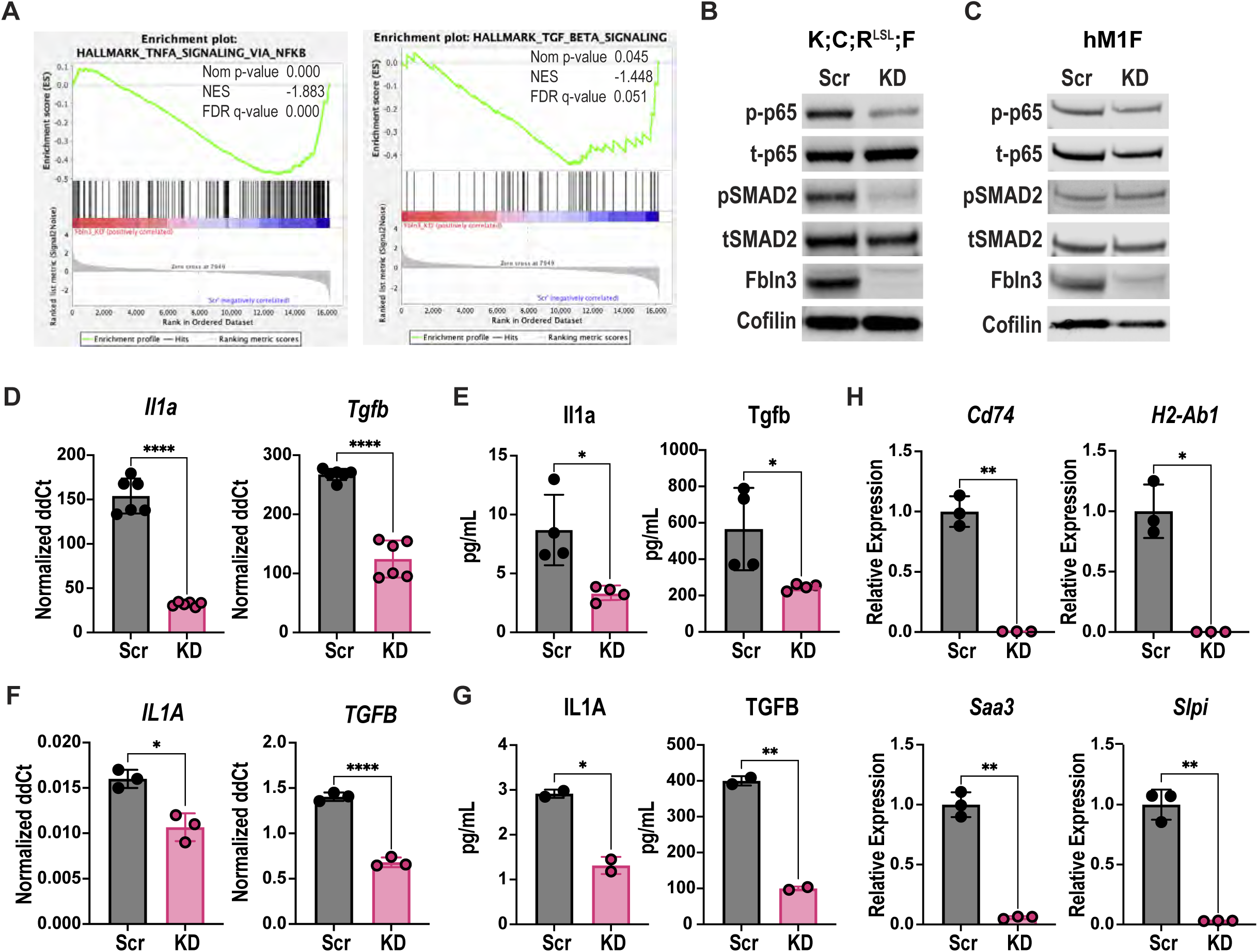
**A)** GSEA comparing Fbln3 knockdown (KD) versus scrambled control (Scr) in CA19-9–expressing K;C;R^LSL^;F organoids. NES, normalized enrichment score; FDR: false discovery rate. **B)** Immunoblot of phosphorylated and total (p/t) p65 (NF-κB), SMAD2, and Fbln3 in CA19-9–expressing K;C;R^LSL^;F organoids with Scr or Fbln3 KD. Cofilin served as a loading control. **C)** Immunoblot of phosphorylated and total (p/t) p65 (NF-κB), SMAD2, and Fbln3 in human patient-derived organoids (hM1F) with Scr or Fbln3 KD. Cofilin served as a loading control. **D)** qRT-PCR measurement of *Il1a* and *Tgfb* expression in CA19-9–expressing K;C;R^LSL^;F organoids with Scr or Fbln3 KD. Biological replicates: n=2, technical replicates: n=3. Statistical analysis by unpaired, two-tailed Welch’s corrected t test. **E)** Protein levels of secreted Il1a and Tgfb in CM from CA19-9–expressing K;C;R^LSL^;F organoids with Scr or Fbln3 KD. Biological replicates: n=2, technical replicates: n=3. Statistical analysis by Mann-Whitney test. **F)** qRT-PCR measurement of *IL1a* and *TGFB* expression in human patient-derived organoids (hM1F) with Scr or Fbln3 KD. Technical replicates: n=3. Statistical analysis by unpaired, two-tailed Welch’s corrected t test. **G)** Protein levels of secreted IL1A and TGFB in CM from human patient-derived organoids (hM1F) with Scr or Fbln3 KD. Technical replicates: n=2. Statistical analysis by unpaired, two-tailed Welch’s corrected t test. **H)** PanMESO cells were treated with CM from CA19-9–expressing K;C;R^LSL^;F organoids with Scr or Fbln3 KD for 72 hours. Cells were harvested and analyzed by qPCR for apCAF markers: Cd74, H2-Ab1 (MHCII), Saa3, and Slpi. *n=3*. Data shown as mean ± SD. Statistical analysis by unpaired, two-tailed Welch’s corrected t test.

To directly assess the functional role of FBLN3 on Il1a and Tgfb production in human PDAC cells, we knocked down FBLN3 in CA19-9^pos^ human PDAC patient-derived organoids (hPDOs) (**Fig. S8A**). Knocking down FBLN3 significantly reduced IL1a and TGFB expression (**Fig. 6F-G**) and suppressed downstream EGFR, STAT3, p65, and SMAD2 signaling pathways (**Fig. 6C, S8B**). Blocking FBLN3 using a fully humanized FBLN3-neutralizing antibody^36^ also decreased downstream EGFR, STAT3, p65, and SMAD2 signaling pathways (**Fig. S8C**).

To determine whether Fbln3-mediated Il1a and Tgfb production functionally contributes to apCAF differentiation, we treated PanMeso cells with CM from control or *Fbln3* KD KCR^LSL^F dox-treated organoids. We found that CM from *Fbln3* KD KCR^LSL^F organoids significantly reduced apCAF differentiation relative to CM from Scr control organoids (**Fig. 6H**). These findings demonstrated that Fbln3-mediated upregulation of Il1a and Tgfb induces apCAF and Treg differentiation.

To test whether Il1a and Tgfb expression is regulated by Egfr hyperactivation, we treated KCR^LSL^F organoids with Dox and the pan-Erbb kinase inhibitor (afatinib). Inhibition of the Egfr pathway in these organoids resulted in a significant reduction in Il1a and Tgfb gene and protein expression (**Fig. S8D-E**). Moreover, treatment of PanMESO cells with CM from afatinib-treated organoids significantly suppressed apCAF differentiation relative to vehicle control (**Fig. S8F**). Afatinib treatment did not prevent recombinant Il1a and Tgfb induced differentiation of PanMeso cells into apCAFs (**Fig. S8G**) Together these data suggest that Fbln3-mediated hyperactivation of Egfr induces Il1a and Tgfb production in CA19-9^pos^ malignant epithelial cells, which in turn promotes apCAF differentiation.

### Fbln3 knockdown affects tumor growth in murine PDAC

To investigate the impact of Fbln3 on the TME *in vivo*, we utilized a genetic approach given the limited efficacy of the Fbln3 blocking antibodies, likely due to their low affinity (data not shown). To accelerate investigation in mice, we generated *Fbln3* KD KPC-derived 2D PDAC cell lines that constitutively express CA19-9. These lines form tumors within weeks, whereas tumor formation of *Kras*-mutant organoids requires >6 months^34^. Fbln3 was efficiently silenced in three independent cell lines (**Fig. S9A**). Across all three, Il1a expression was significantly reduced with Fbln3 loss, while Tgfb expression was decreased in two of the three lines (**Fig. S9B**). A similar trend of reduced Il1a and Tgfb protein levels was observed (**Fig. S9C**). We observed no differences in growth of these cell lines *in vitro* using either full- or reduced-serum conditions (**Fig. S9D**).

To examine the contribution of Fbln3 on the TME *in vivo*, we orthotopically transplanted CA19-9^pos^ control or *Fbln3* KD KPC 2D cells into syngeneic C57Bl/6J mice and confirmed loss of Fbln3 levels (**Fig. 7A, S10A**). *Fbln3* KD tumors exhibited significantly slower growth kinetics than controls (**Fig. 7B, S10B**), and tumor weight, volume, and size were all significantly decreased (**Fig. 7C-D, S10C-D**). Despite these differences in tumor burden, Ki67 and Cc3 IHC indicated no significant changes in tumor cell proliferation or apoptosis at endpoint (**Fig. S10E-F**). Additionally, pEgfr levels exhibited a trend towards lower levels in *Fbln3* KD tumors relative to controls (**Fig. S10E-F**). Immunofluorescence analysis revealed significant reductions in total CAFs and apCAFs in *Fbln3* KD tumors (**Fig. 7E**). Although Treg abundance remained unchanged (**Fig. 7F, S10G**), we found that Cd8^+^ T cell infiltration was significantly and substantially increased in Fbln3 KD tumors (**Fig. 7G, S10H**). These data show that Fbln3 loss leads to accumulation of Cd8^+^ T cells.

**Figure 7.**
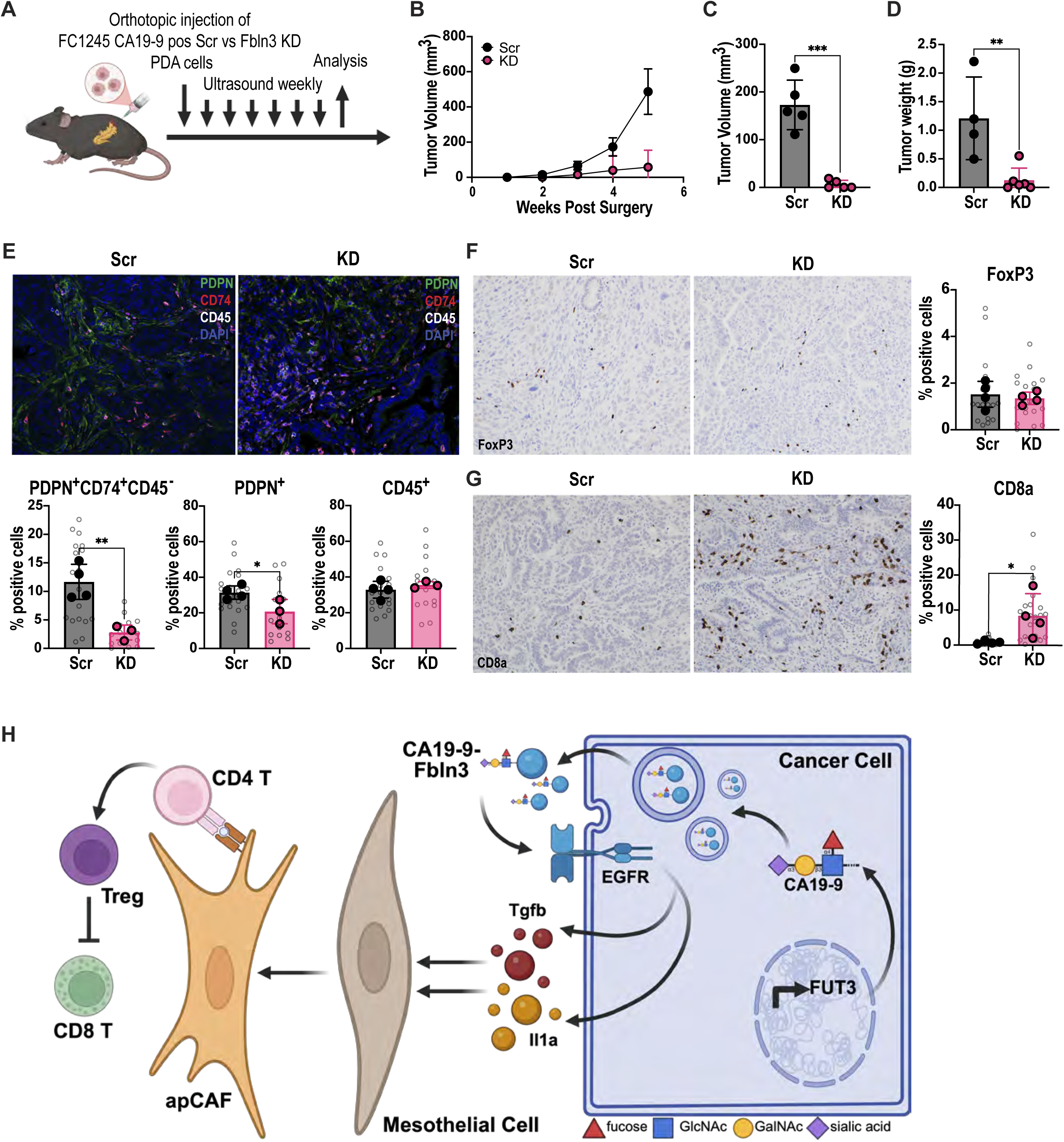
**A)** Schematic of orthotopic transplantation mouse study **B)** Tumor volume over time following orthotopic transplantation of FC1245 cells with Scr or Fbln3 KD. **C)** Tumor volume comparison 5 weeks post-injection of FC1245 Scr vs Fbln3 KD cells. Statistical analysis by unpaired, two-tailed t test. **D)** Tumor weight comparison 5 weeks post-injection of FC1245 Scr vs Fbln3 KD cells. Statistical analysis by unpaired, two-tailed t test. **E)** IF and quantification of Pdpn^+^Cd74^+^Cd45^-^ (apCAF), Pdpn^+^ (pan-CAFs), and Cd45^+^ (pan-immune cells) in orthotopic tumors. Bar graphs represent the mean, and error bars represent the standard deviation. n=4, 3. Each grey circle represents one FOV. Each solid data point represents the average of all FOVs/mouse (20X magnification). Statistical analysis by unpaired, two-tailed t test. **F)** IHC and quantification of FoxP3 (Tregs) in orthotopic tumors. Bar graphs represent the mean, and error bars represent the standard deviation. n=4. Each grey circle represents one FOV. Each solid data point represents the average of all FOVs/mouse (20X magnification). Statistical analysis by unpaired, nonparametric two-tailed Mann-Whitney test. **G)** Cd8a (Cd8^+^ T cells) in orthotopic tumors. Bar graphs represent the mean, and error bars represent the standard deviation. n=4. Each grey circle represents one FOV. Each solid data point represents the average of all FOVs/mouse (20X magnification). Statistical analysis by unpaired, nonparametric two-tailed Mann-Whitney test. **H)** Schematic summary of paper.

To assess the contribution of anti-tumor immunity to Fbln3-mediated tumor growth, we orthotopically transplanted CA19-9^pos^ control or *Fbln3* KD KPC 2D PDAC cells into immunodeficient mice and evaluated tumor burden. In the absence of functional T, B, and NK cells, there was no significant difference in tumor size (**Fig. S10I-J**). These data suggest that Fbln3 accelerates PDAC progression by restricting Cd8^+^ T cell infiltration, most likely through the action of apCAF-mediated Treg differentiation, leading to an immunosuppressive TME.

Altogether, we show that CA19-9 induces hyperactivation of Egfr in the ductal compartment through Fbln3 (**Fig. 7H**). This leads to increased production of Il1a and Tgfb, thereby driving apCAF differentiation. apCAFs directly engage with Cd4^+^ T cells, causing Treg differentiation and reduced Cd8^+^ T cell infiltration.

## DISCUSSION

CA19-9 has traditionally been viewed as a passive biomarker of disease burden. Our findings instead reveal CA19-9 is an active regulator of the PDAC TME. Elevated CA19-9 initiates a signaling cascade in which CA19-9-modified Fbln3 promotes EGFR-dependent production of Il1a and Tgfb by malignant epithelial cells. These cytokines induce mesothelial differentiation into apCAFs, driving Treg expansion and immunosuppression.

Therapeutic targeting of CA19-9 has been evaluated in patients with metastatic PDAC who progressed on chemotherapy. Antibody monotherapy produced stable disease in 20% of patients (>90 days), and combination treatment with chemotherapy produced partial responses in 27% of patients lasting more than 10 months. These responses are notable in advanced -line metastatic PDAC. However, efficacy was restricted to patients with serum CA19-9 levels below 2,500 U/mL. Higher levels were associated with increased antibody clearance and reduced therapeutic activity. This pharmacokinetic limitation is clinically relevant because patients with metastatic PDAC often have CA19-9 levels exceeding 10,000 U/mL. These data identify a key limitation of directly targeting CA19-9.

Our dissection of CA19-9-regulated signaling networks identifies alternative therapeutic opportunities that circumvent anti-CA19-9 antibody clearance limitations. Il1a and Tgfb regulate CAF lineage specification, exerting opposing effects on iCAF and myCAF states^19^. Individual targeting of these cytokines in combination with standard chemotherapy regimens such as gemcitabine and nab-paclitaxel but did not improve outcomes^48,49^. Combined targeting of IL1A and TGFB may warrant further exploration. Targeting Fbln3 represents a promising strategy that relieves immunosuppressive pressure within the PDAC TME and increases Cd8^+^ T cell infiltration. Unlike pan-ErbB inhibition, which is associated with grade 3 toxicities in 26-49% of patients^50^, Fbln3 targeting could selectively attenuate oncogenic EGFR signaling. Because CA19-9 modification enhances the pathogenic activity of Fbln3, antibodies or pharmacologic agents that specifically recognize CA19-9-modified Fbln3 could provide strong tumor selectivity. Defining the structural consequences of CA19-9 modification, including effects on Fbln3 affinity for EGFR, localization, stability, and protein interactions, will be important for therapeutic development. More broadly, the observation that CA19-9 modification alters Fbln3 signaling raises the possibility that glycan modifications regulate the signaling properties of other glycoproteins, potentially revealing additional therapeutic targets.

PDAC is characterized by a multi-layered immunosuppressive microenvironment, employing redundant mechanisms including myeloid-derived suppressor cells, tumor-associated macrophages, T cell exhaustion, and desmoplasia. Our findings position the CA19-9/Fbln3/apCAF/Treg axis as a critical and potentially foundational layer to this ecosystem. Several lines of evidence suggest this is not merely an additive mechanism but may actively coordinate broader immunosuppression: Tregs recruit suppressive myeloid populations and inhibit effector T cells, apCAFs promote stromal barriers to T cell infiltration, and Il1a and Tgfb exert pleiotropic effects across multiple cell types^13,17,19^. The improved prognosis of CA19-9-negative PDAC patients further suggests this axis is functionally non-redundant. However, it remains unclear whether the apCAF and Treg axis acts as an early initiating event or operates in parallel with other pathways through reinforcing feedback loops. Understanding these interactions will be essential for designing effective combination immunotherapies.

These findings establish a framework in which glycosylation orchestrates multicellular signaling networks that shape tumor ecosystems. Capturing these effects requires models that recapitulate human glycan biology. Most genetically engineered mouse models of PDAC lack CA19-9 expression and may therefore fail to reproduce the full immunosuppressive landscape of human disease. This limitation may contribute to the poor clinical translation of immunotherapies developed in preclinical models. Our results highlight glycan-dependent signaling as a previously underappreciated mechanism of tumor immune regulation. By linking glycobiology to defined immunosuppressive circuits, this work suggests that aberrant glycosylation can actively instruct tumor-stromal-immune crosstalk. Systematic interrogation of glycan-regulated signaling pathways may therefore uncover an expanded class of immunotherapeutic targets across malignancies characterized by dysregulated glycosylation.

## Author contributions

JH, HS, and DDE designed research studies. JH, HS, SO, CSK, KLP, EJ, LGR, JZ, YS, WJ, YD, JL, CRB, KC, MB, EK, KK, AHM, MS, AR, and SRO conducted experiments and acquired data. JZ, WJ, TGO performed bioinformatic analyses. TH, MD, RE, JZ, TGO, YZ, AML, HT, and SMK provided experimental guidance and reagents. JH, HS, and DDE wrote the manuscript with input from all authors.

## DATA AVAILABILITY STATEMENT

All data associated with this study are available in the article or the Supporting Data Values file. Mouse bulk and single cell RNA-seq data generated in this study will be deposited in the Gene Expression Omnibus (GEO) upon publication.

## Declaration of generative AI and AI-assisted technologies in the manuscript preparation process

During the preparation of this work the authors used ChatGPT in order to edit grammar and syntax. After using this tool/service, the authors reviewed and edited the content as needed and take full responsibility for the content of the published article.

## ACKNOWLEDGMENTS

PanMeso cells were kindly provided by Dr. Huocong Huang and Rolf Brekken. The authors would like to thank all the members and alumni of the Engle laboratory, the Animal Resources Department (ARD), Daniela Boassa, Sammy Weiser Novak and Elsie Quansah at the Biophotonics Core Facility (BPHO), Flow Cytometry Core Facility (FCCF), and the Integrated Genomics and Bioinformatics Core (IGC) of the Salk Institute.

## FIGURE LEGENDS

**Supplementary Figure 1.**
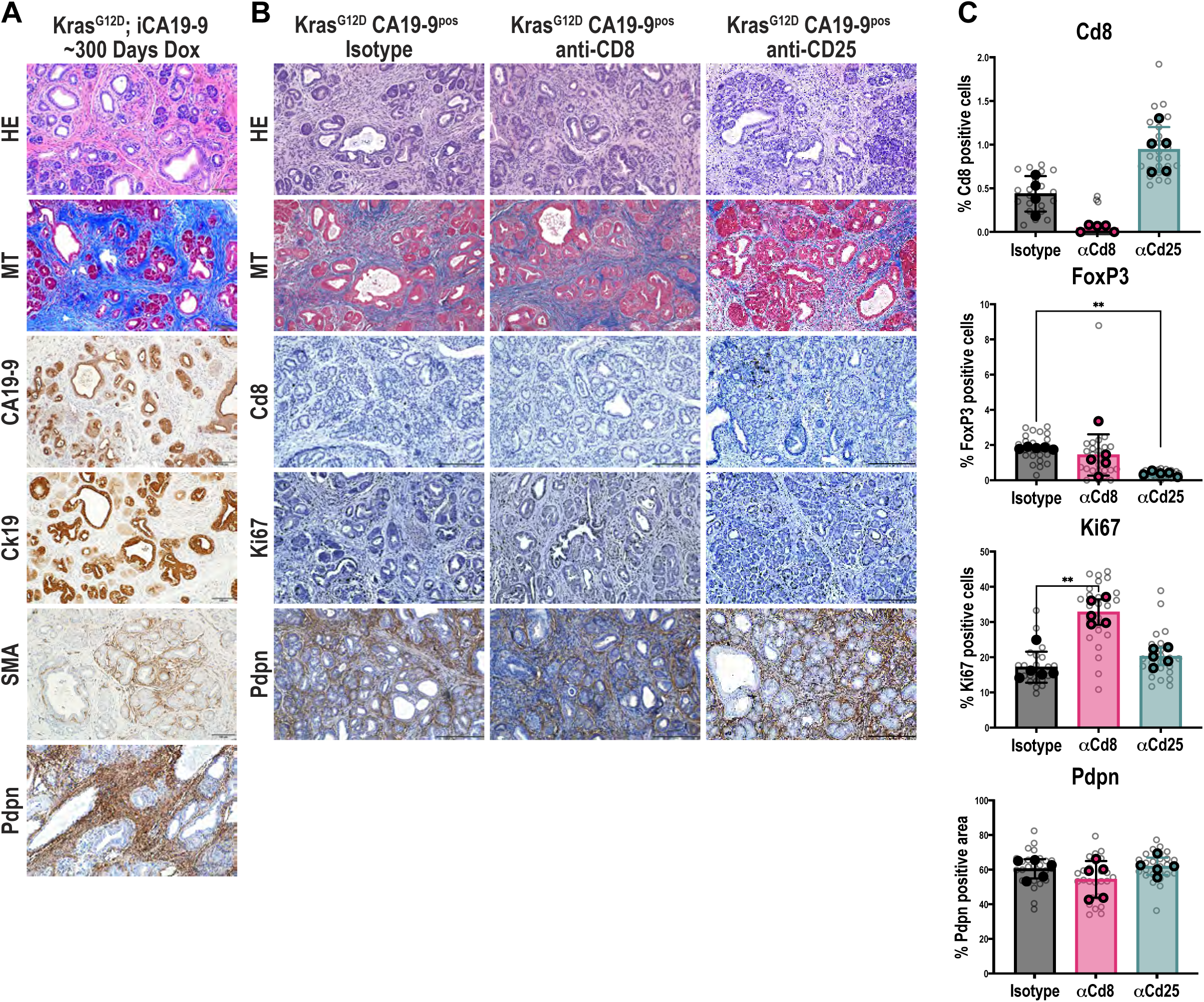
**A)** Histologic evaluation of KCR^LSL^F pancreata at humane endpoint after treatment with Dox by HE and MT staining. CA19-9, Ki67, smooth muscle actin (aSMA), and pdpn expression evaluated by IHC. Scale bars = 100um. **B)** Histologic evaluation of KCR^LSL^F pancreata after treatment of Dox and anti-Cd8 and anti-Cd25 antibody blockade compared to isotype control by HE and MT staining. Cd8, Ki67, and Pdpn expression evaluated by IHC. Scale bars = 100um. **C)** Cd8, FoxP3, Ki67 histological quantification as a percentage of total cells in single FOVs and Pdpn quantification as a percentage of total tissue area in single FOVs in KCR^LSL^F+Dox pancreata and anti-Cd8 and anti-Cd25 antibody blockade compared to isotype control. Bar graphs represent the mean, and error bars represent the standard deviation. n=5. Each grey circle represents one FOV. Each solid data point represents the average of all FOVs/mouse. Statistical analysis by Kruskal-Wallis test.

**Supplementary Figure 2.**
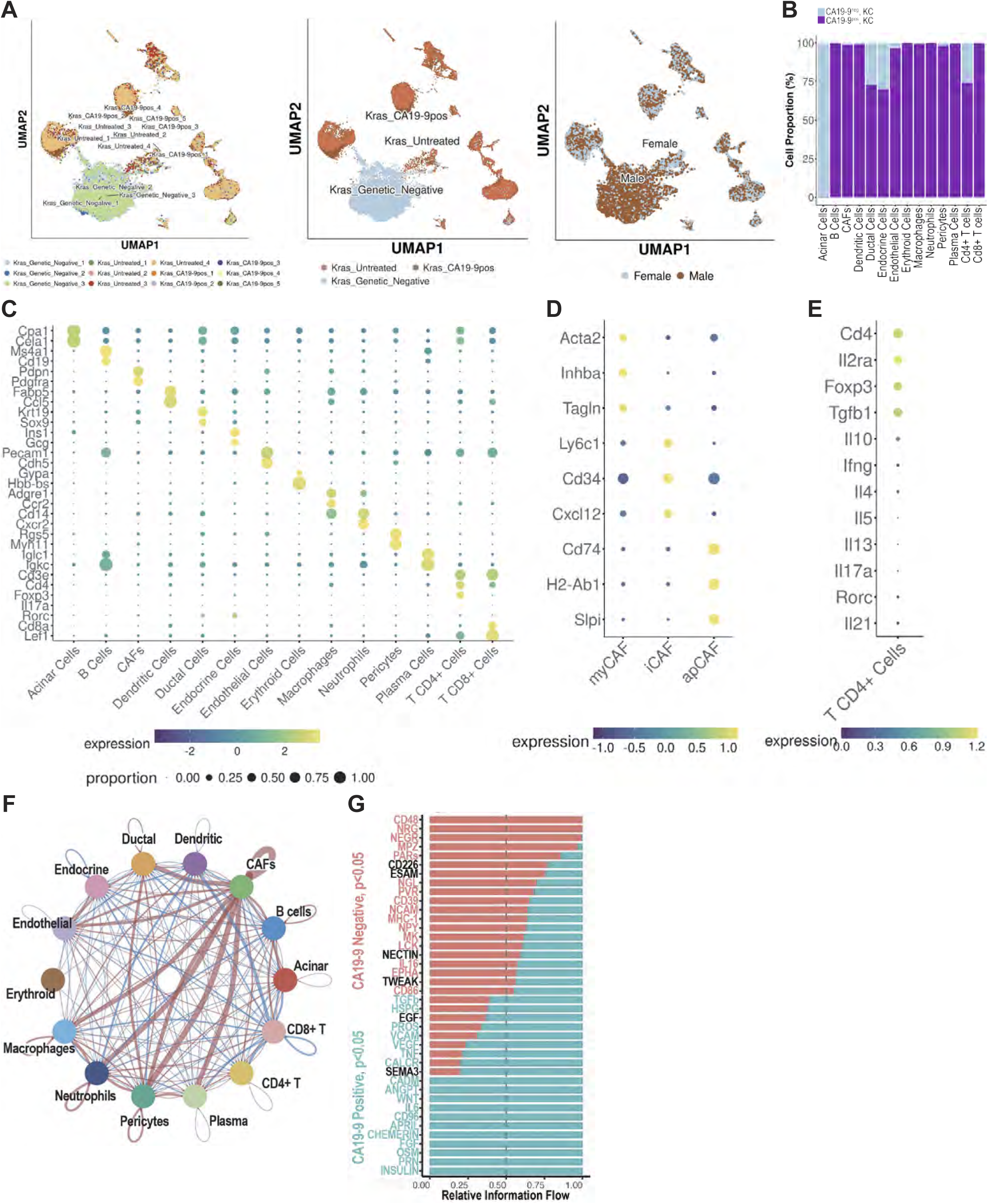
**A)** UMAP projection of distribution of single cells per replicate samples of KC, KCR^LSL^F treated and untreated mouse pancreata (left) and pooled arms (middle). UMAP projection of mouse sex of all single cell samples (right). **B)** Cell proportion graph from scRNAseq showing percentage cell proportion of all cell type clusters from UMAP projections. **C)** Gene expression plot from scRNAseq of all cell types and cell type defining genes. **D)** Gene expression plot from scRNAseq of CAF subtypes and cell type defining genes. **E)** Gene expression plot from scRNAseq of Cd4^+^ T cells and cell type defining genes. **F)** CellChat projection of cell type clusters from UMAP. Red and blue lines denote upregulation and downregulation of signaling, respectively. Thickness of line denotes magnitude of change. **G)** Relative information flow graph from CellChat displaying top 20 genes in CA19-9^pos^ and CA19-9^neg^ pancreata

**Supplementary Figure 3.**
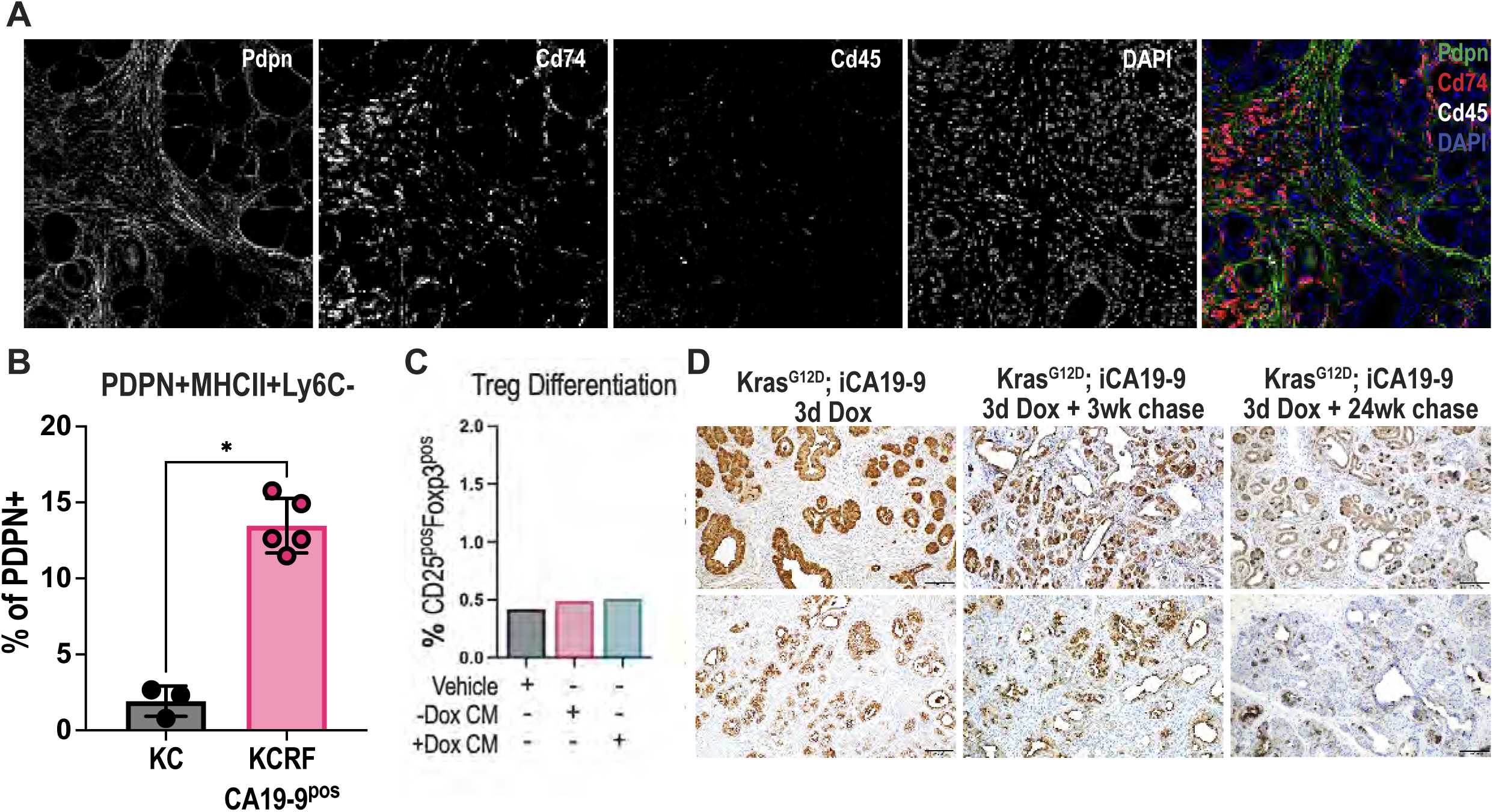
**A)** Single channels of apCAF IF (Pdpn^+^Cd74^+^Cd45^-^) in KCR^LSL^F mice treated with Dox for 3 days. **B)** Flow cytometry quantification of KC and KCR^LSL^F pancreata treated with Dox. apCAFs (PDPN^+^MHCII^+^Ly6C^-^) were quantified as a percentage of PDPN^+^ cells from single cell dissociated pancreata. Bar graphs represent the mean, and error bars represent the standard deviation. n=5. Solid dots represent individual mice. Statistical analysis by unpaired, two-tailed Welch’s corrected t test. **C)** Flow cytometry quantification of Treg (Cd25+FoxP3+) differentiation of OT-II Naive Cd4^+^ T cells cultured in KCR^LSL^F organoid CA19-9 positive and negative CM. n=1. Statistical analysis by ordinary one-way ANOVA. **D)** Histologic evaluation of KCR^LSL^F pancreata after treatment with Dox for 72 hours and subsequent 3-week and 24-week chase by GFP and CA19-9 IHC staining. Scale bars = 100um.

**Supplementary Figure 4.**
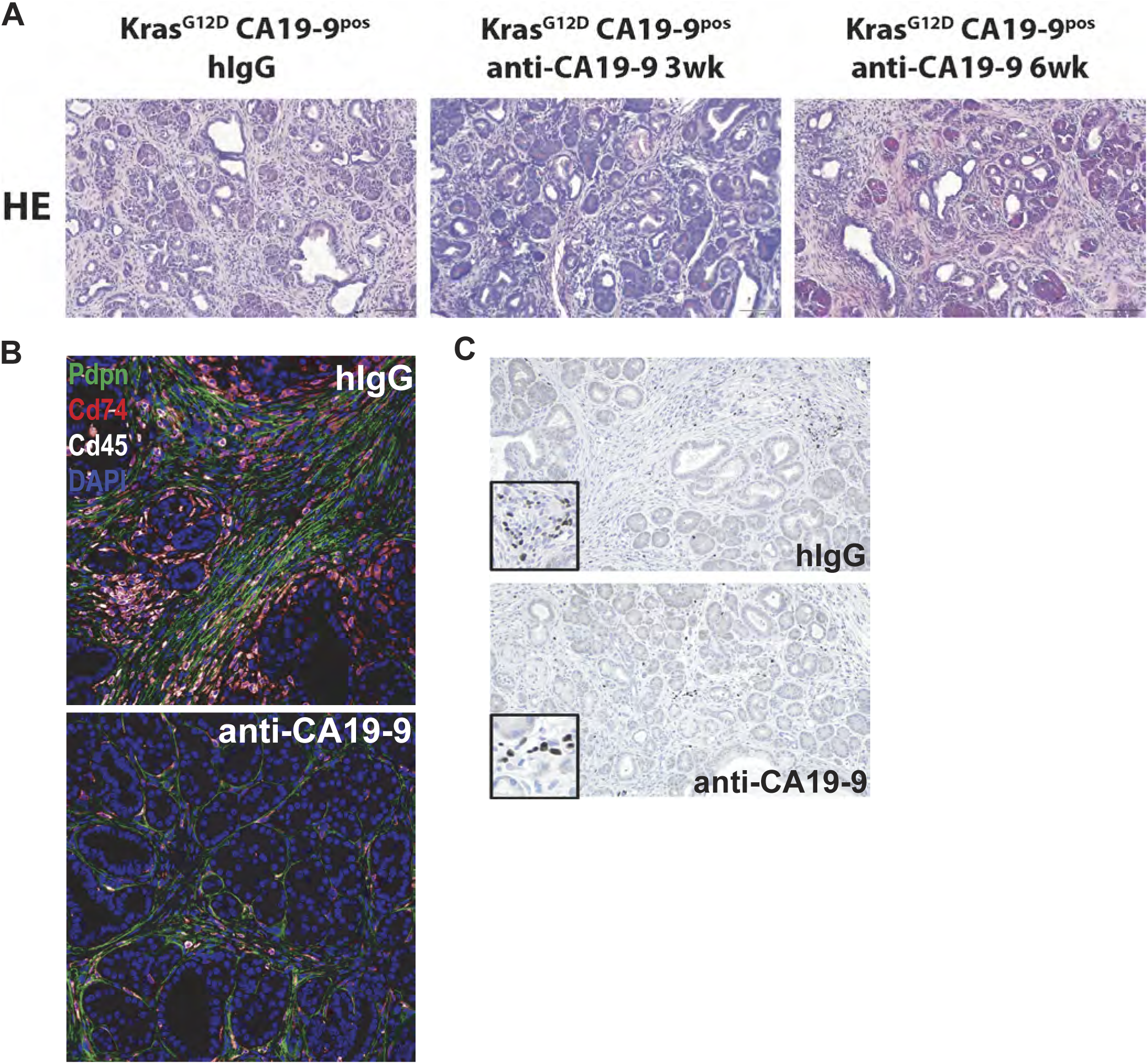
**A)** Histologic evaluation by HE staining of KCR^LSL^F pancreata after the treatment with Dox followed by 3 and 6 weeks of weekly injections of anti-CA19-9 vs isotype control. Scale bars = 100 um. **B)** Representative apCAF IF image in the anti-CA19-9 in vivo study. **C)** Representative FoxP3 IHC image in the anti-CA19-9 in vivo study.

**Supplementary Figure 5.**
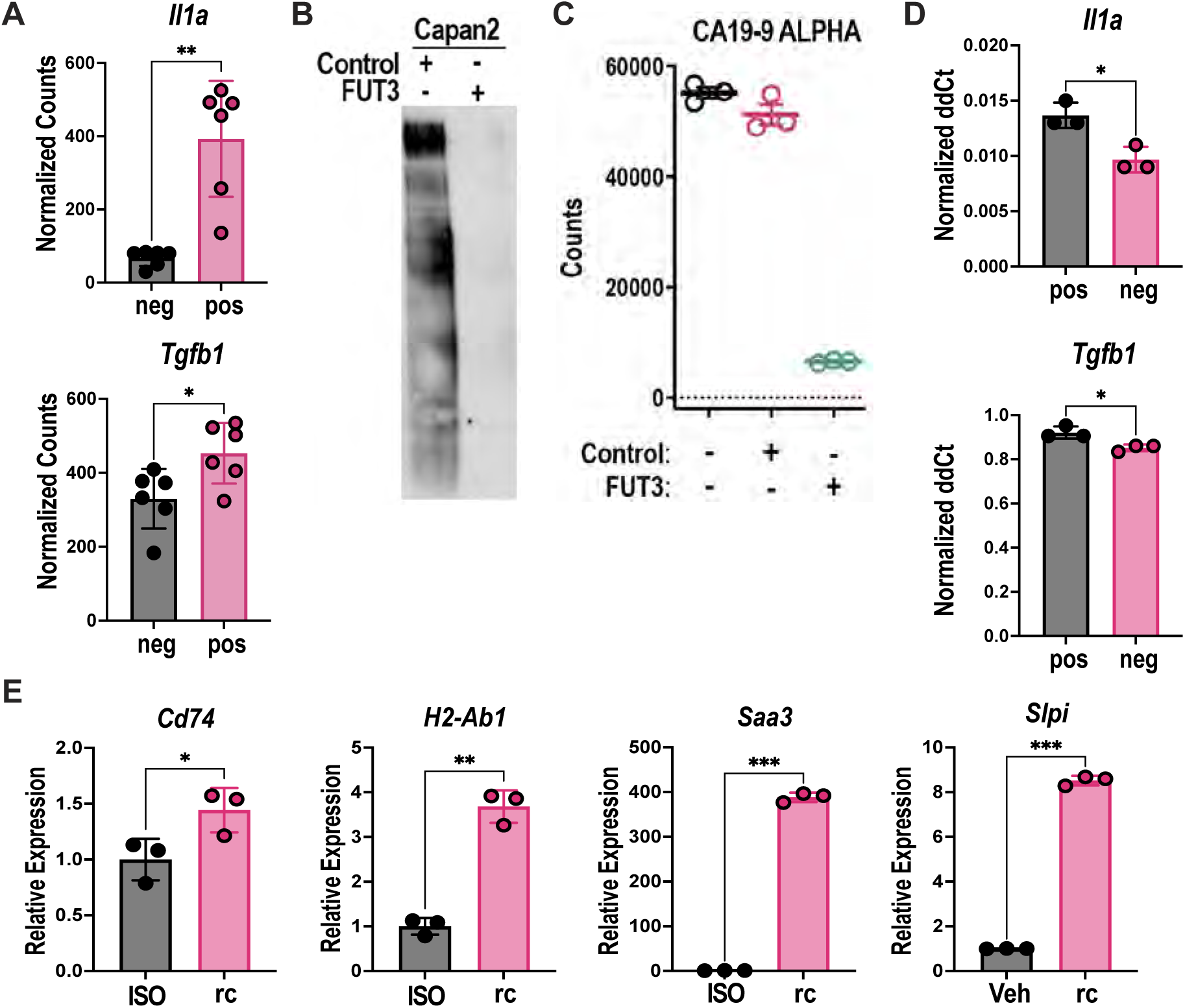
**A)** RNA-seq analysis of *Il1a* and *Tgfb* gene expression levels. Biological replicates: n=2. Technical replicates: n=3. Data shown as mean ± SD. Statistical analysis by unpaired, two-tailed Welch’s corrected t test. **B)** Immunoblot of CA19-9 in Capan2 cells with FUT3 and Rosa26 locus control knockout. **C)** ALPHA (Amplified Luminescent Proximity Homogeneous Assay) bead-based quantification of CA19-9 in conditioned media from Capan2 cells with FUT3 and Rosa26 locus control knockout. **D)** qRT-PCR of *IL1a* and *TGFb* in Capan2 cells with FUT3 KO or Rosa26 locus control. Statistical analysis by unpaired, two-tailed Welch’s corrected t test. **E)** PanMESO cells were treated with CM from K;C;R^LSL^;F organoids pre-treated with both Dox and IL1A and TGFB recombinant proteins (rc) or IgG isotype controls (ISO) for 72 hours. Cells were harvested and analyzed by qPCR for apCAF markers: Cd74, H2-Ab1 (MHCII), Saa3, and Slpi. *n=3*. Data shown as mean ± SD. Statistical analysis by unpaired, two-tailed Welch’s corrected t test.

**Supplementary Figure 6.**
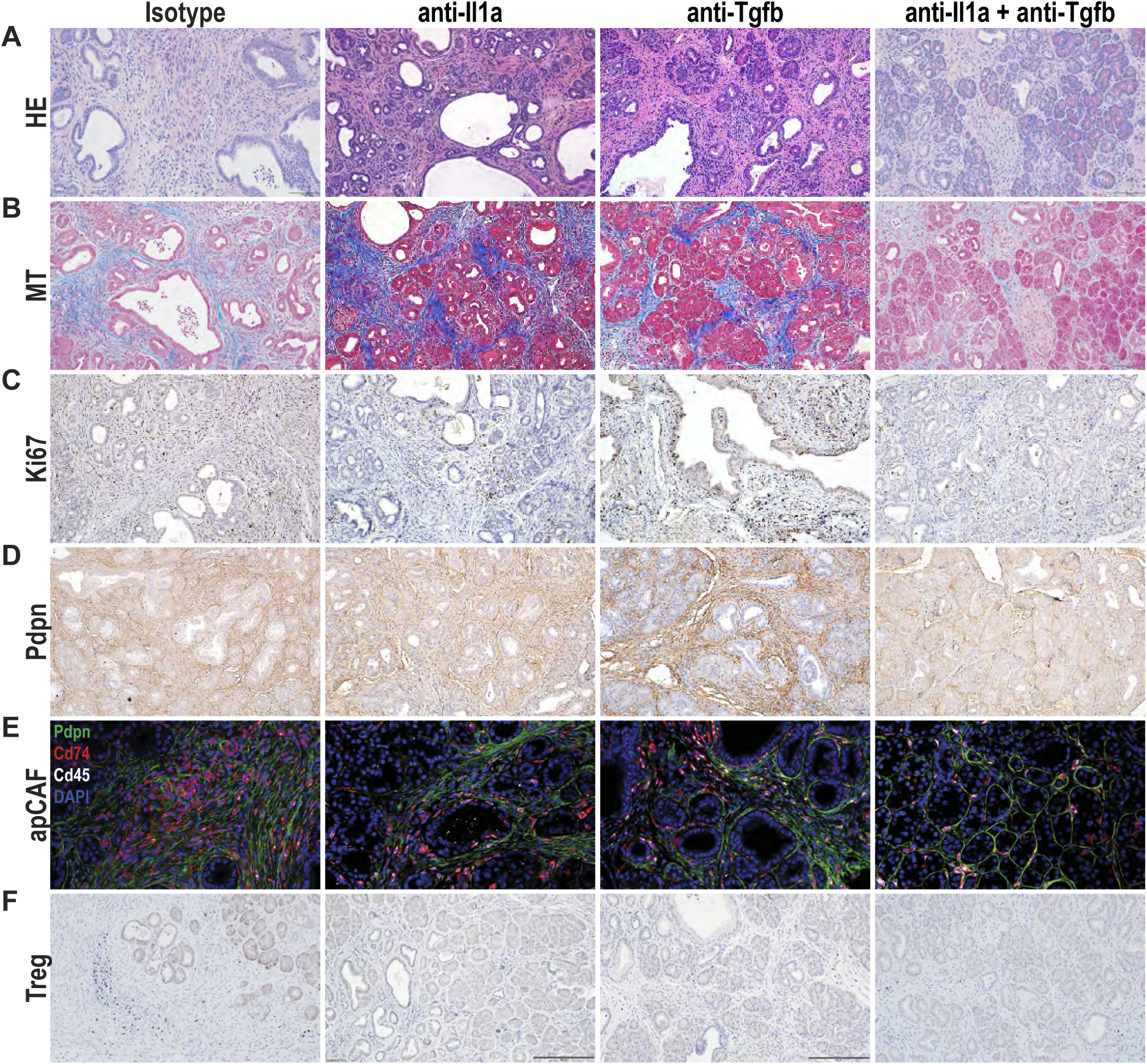
**A)** Representative H&E of KCR^LSL^F pancreata after the treatment with Dox with anti-Il1a, -Tgfb antibody blockade, individually and in combination, compared to isotype control (n=5, 5, 5, 10 respectively). **B)** Representative MT of KCR^LSL^F pancreata after the treatment with Dox with anti-Il1a, -Tgfb antibody blockade, individually and in combination, compared to isotype control (n=5, 5, 5, 10 respectively). **C)** Representative Ki67 IHC of KCR^LSL^F pancreata after the treatment with Dox with anti-Il1a, - Tgfb antibody blockade, individually and in combination, compared to isotype control (n=5, 5, 5, 10 respectively). **D)** Representative Pdpn IHC of FOVs in KCR^LSL^F pancreata after the treatment with Dox with anti-Il1a, -Tgfb antibody blockade, individually and in combination, compared to isotype control (n=5, 5, 5, 10 respectively). **E)** Representative apCAF IF of KCR^LSL^F pancreata after the treatment with Dox with anti-Il1a, - Tgfb antibody blockade, individually and in combination, compared to isotype control (n=5, 5, 5, 10 respectively). **F)** Representative FoxP3 IHC of KCR^LSL^F pancreata after the treatment with Dox with anti-IL1a, -TGFb antibody blockade, individually and in combination, compared to isotype control (n=5, 5, 5, 10 respectively).

**Supplementary Figure 7.**
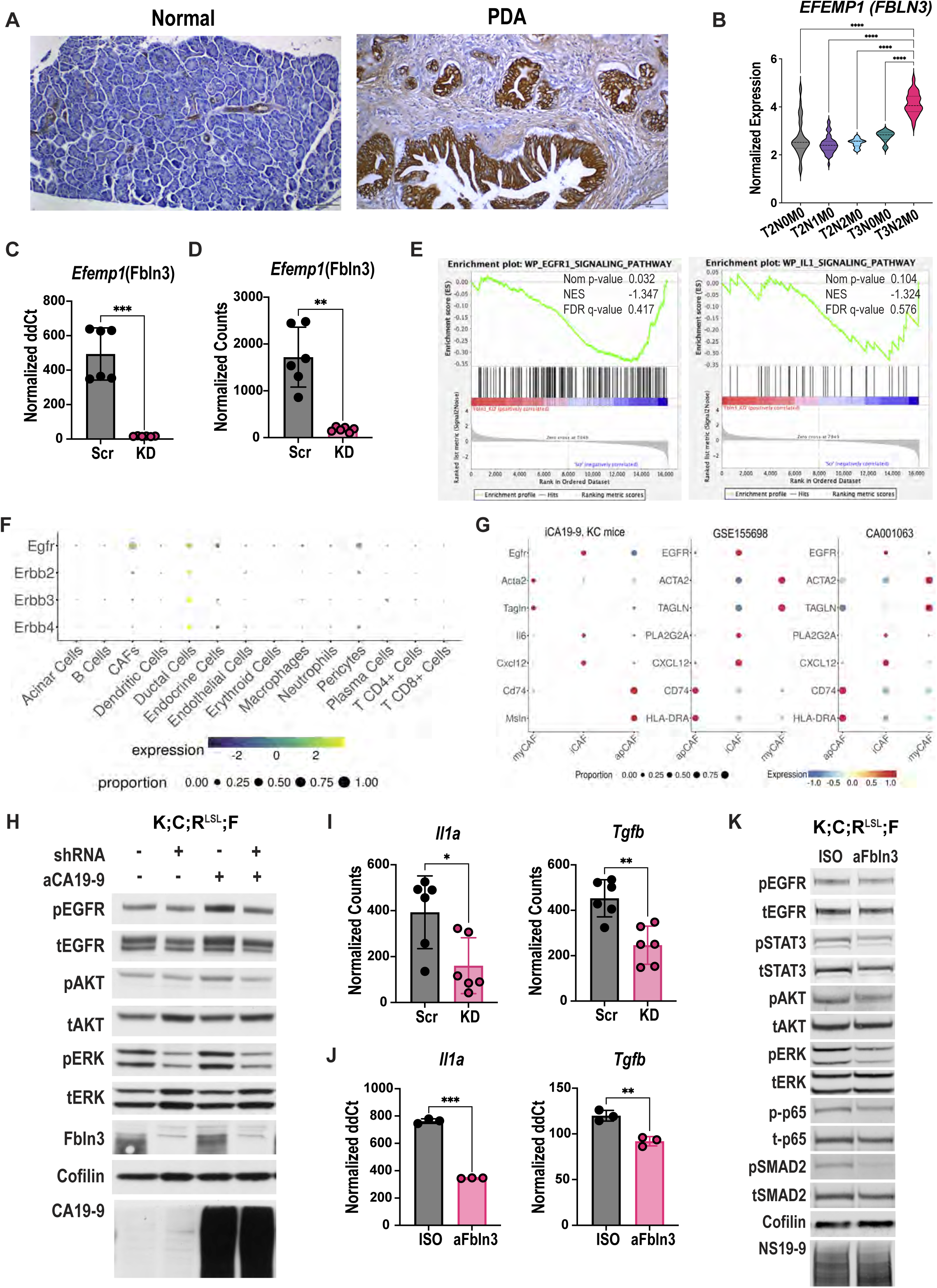
**A)** IHC of Fbln3 in normal human pancreas and PDAC tissues. **B)** Expression levels of *Efemp1* (Fbln3) across PDAC stages using Nanostring GeoMx spatial transcriptomics data^13^. T indicates tumor size/stage (1-4), N denotes lymph node involvement (0-3), and M indicates metastatic status (M0/M1). **C)** qRT-PCR of Fbln3 gene expression in Scr or KD lines. Biological replicates: n=2. Technical replicates: n=3. Statistical analysis by unpaired, two-tailed Welch’s corrected t test. **D)** RNA-seq quantification of Fbln3 gene expression in Scr or KD lines. Biological replicates: n=2. Technical replicates: n=3. Statistical analysis by unpaired, two-tailed Welch’s corrected t test. **E)** GSEA comparing Fbln3 KD versus Scr control in CA19-9–expressing K;C;R^LSL^;F organoids. NES, normalized enrichment score; FDR: false discovery rate. **F)** scRNA-seq analysis of Erbb family gene expression across cell types in the K;C;R^LSL^;F pancreata after 3 day Dox treatment. **G)** scRNAseq analyses of CAF subtype defining genes and EGFR in K;C;R^LSL^;F pancreata after 3 day Dox treatment as well as 2 different human scRNA-seq public datasets (GSE155698, CA001063). **H)** Immunoblot analysis of p/t EGFR, AKT, and ERK in mouse CA19-9 inducible K;C;R^LSL^;F organoids with Scr of Fbln3 KD. Organoids were treated or untreated with Dox. Cofilin served as a loading control. **I)** RNA transcript measurement of *Il1a*, and *Tgfb* from RNA-seq analysis. Statistical analysis by unpaired, two-tailed Welch’s corrected t test. **J)** qPCR measurement of *Il1a* and *Tgfb* expression in CA19-9 expressing, Dox treated K;C;R^LSL^;F organoids treated with mIgG or Fbln3-blocking antibody (4ug/ml). Statistical analysis by unpaired, two-tailed Welch’s corrected t test. **K)** Immunoblot analysis of p/t EGFR, STAT3, ERK, p65 (NF-kB), and SMAD2 in mouse CA19-9 expressing, Dox treated K;C;R^LSL^;F organoids treated with mIgG or mAb3-5 (aFbln3) (4ug/mL) for 48 hours.

**Supplementary Figure 8.**
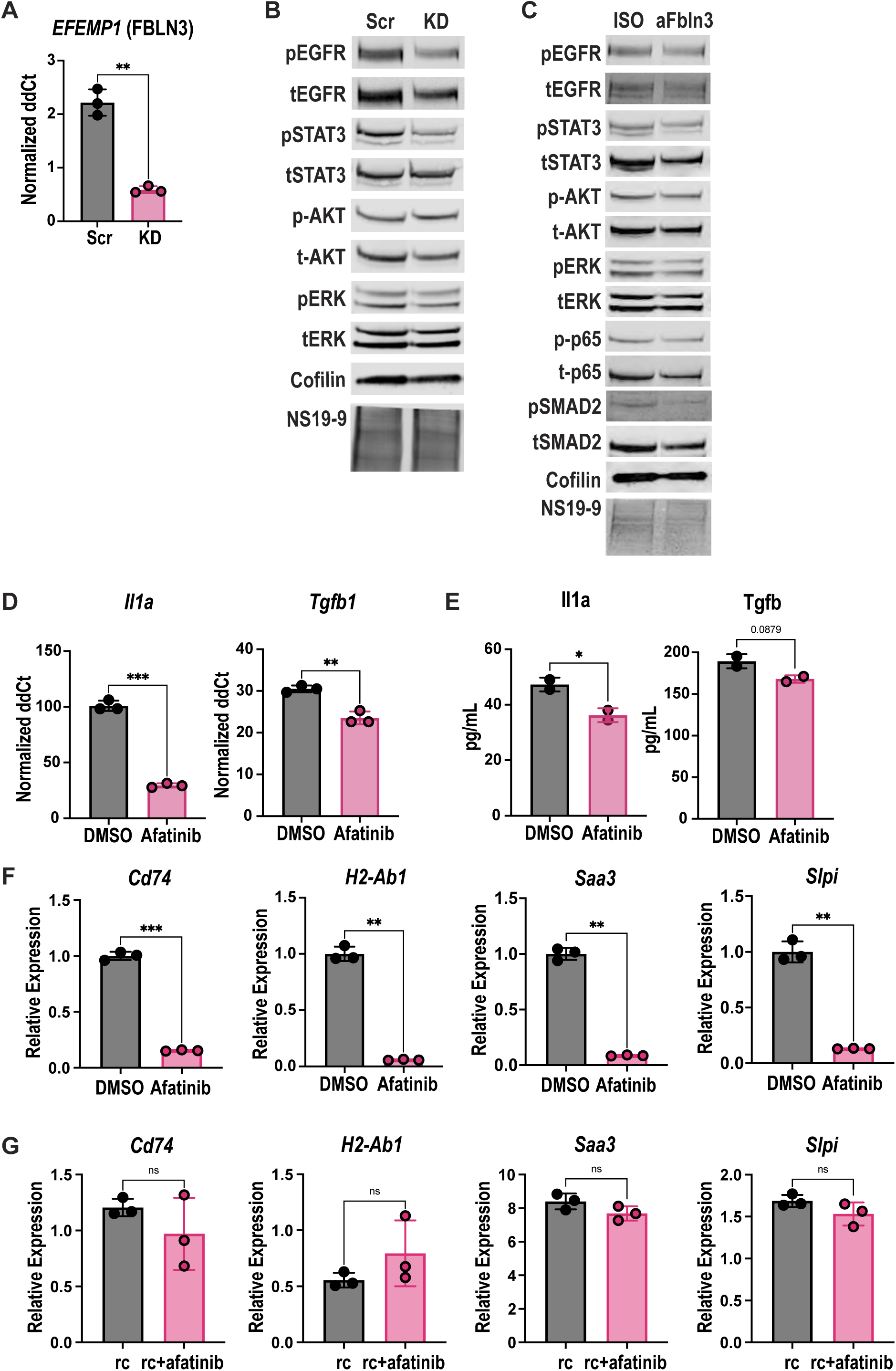
**A)** qPCR measurement of EFEMP1 (FBLN3) knockdown efficiency in human patient-derived organoid. Statistical analysis by unpaired, two-tailed Welch’s corrected t test. **B)** Immunoblot analysis of p/t EGFR, AKT, and ERK in human patient-derived organoids with Scr of Fbln3 KD. NS19-9 indicates CA19-9. Cofilin served as a loading control. **C)** Immunoblot analysis of p/t EGFR, STAT3, ERK, p65 (NF-kB), and SMAD2 in hPDOs treated with hIgG1 or hFbln3-blocking antibody (25ug/mL) for 48 hours. **D)** qPCR analysis of Il1a and Tgfb1 in CA19-9–expressing K;C;R^LSL^;F organoids treated with DMSO or Afatinib (60nM) for 48 hours. Statistical analysis by unpaired, two-tailed Welch’s corrected t test. **E)** Secreted protein levels of IL1A and TGFB in CM from CA19-9–expressing K;C;R^LSL^;F organoids treated with DMSO or Afatinib (60nM) for 48hours. Statistical analysis by unpaired, two-tailed t test. **F)** PanMESO cells treated with CM from DMSO- or Afatinib-pretreated CA19-9–expressing K;C;R^LSL^;F organoids for 72hours. Cells were harvested and analyzed by qPCR for apCAF markers: Cd74, H2-Ab1 (MHCII), Saa3, and Slpi. *n=3*. Data shown as mean ± SD. Statistical analysis by unpaired, two-tailed Welch’s corrected t test. **G)** PanMESO cells treated with CM from IL1A and TGFB recombinant proteins (rc)- or IL1A and TGFB recombinant proteins with Afatinib-pretreated CA19-9–expressing K;C;R^LSL^;F organoids for 72hours. Cells were harvested and analyzed by qPCR for apCAF markers: Cd74, H2-Ab1 (MHCII), Saa3, and Slpi. *n=3*. Data shown as mean ± SD. Statistical analysis by unpaired, two-tailed Welch’s corrected t test.

**Supplementary Figure 9.**
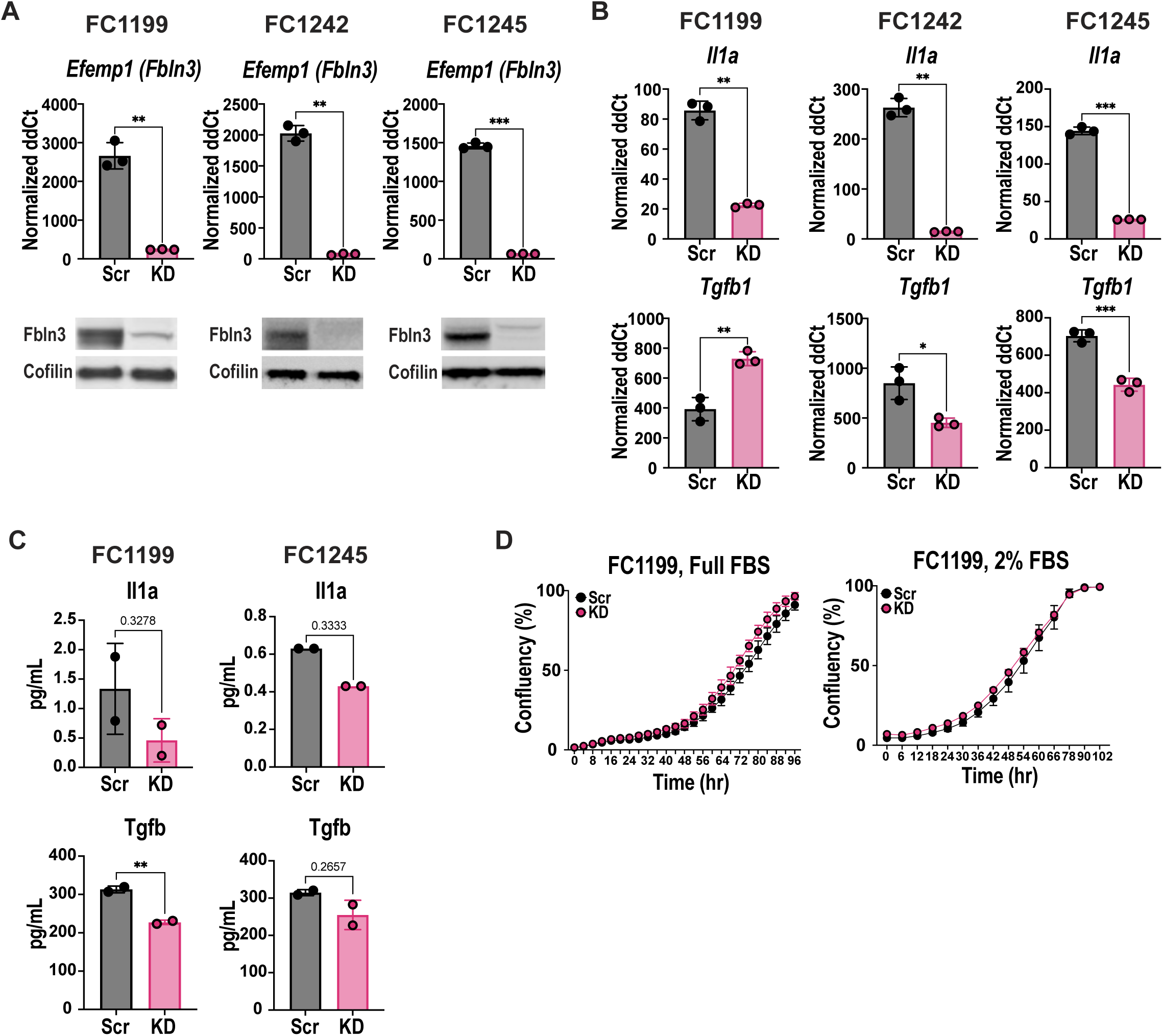
**A)** qPCR and immunoblot analysis of Efemp1 (Fbln3) KD efficiency in CA19-9–expressing KPC-derived PDAC monolayer cell lines (FC1245, FC1199, FC1242). Statistical analysis by unpaired, two-tailed Welch’s corrected t test. **B)** qPCR of Il1a and Tgfb1 expression in Scr vs Fbln3 KD CA19-9-expressing KPC-derived PDAC monolayer cell lines (FC1245, FC1199, FC1242). Statistical analysis by unpaired, two-tailed Welch’s corrected t test. **C)** Protein levels of secreted Il1a and Tgfb in CM from FC1199 and FC1245 with Scr or Fbln3 KD. Technical replicates: n=3. Statistical analysis by unpaired, nonparametric two-tailed Mann-Whitney test. **D)** Cell growth curve of FC1199 cells with Scr of Fbln3 KD in full serum (10% FBS) and low serum (2% FBS) conditions.

**Supplementary Figure 10.**
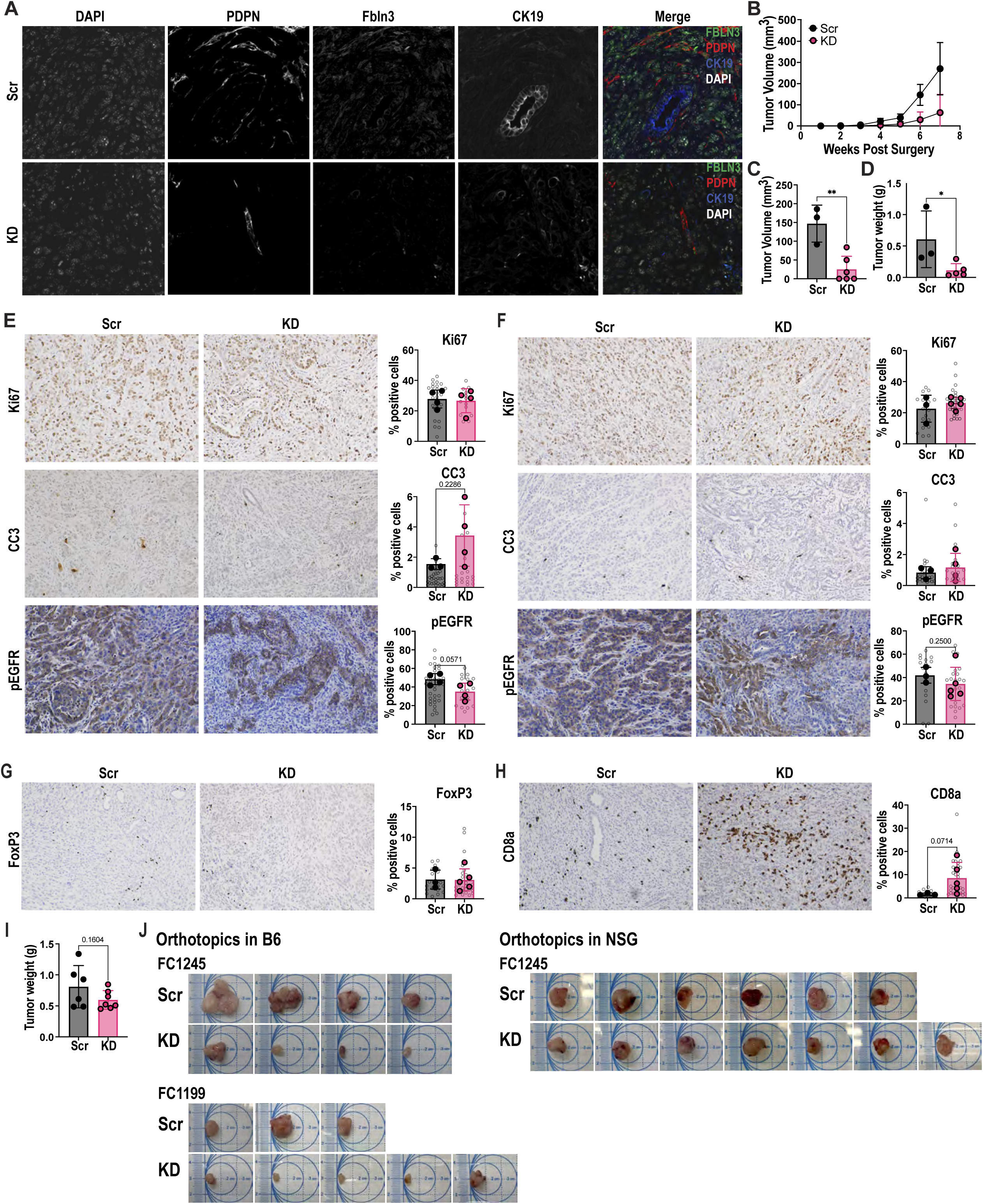
**A)** IF of Fbln3 in FC1245 orthotopic tumors (Scr vs Fbln3 KD) (40X magnification). **B)** Tumor volume over time after orthotopic transplantation of FC1199 Scr vs Fbln3 KD cells. **C)** Tumor volume comparison 7 weeks post-injection of FC1199 Scr vs Fbln3 KD cells. Statistical analysis by unpaired, two-tailed t test. **D)** Tumor weight comparison 7 weeks post-injection of FC1199 Scr vs Fbln3 KD cells. Statistical analysis by unpaired, two-tailed t test. **E)** IHC of Ki67, CC3, and pEGFR in FC1245 orthotopic tumors. Bar graphs represent the mean, and error bars represent the standard deviation. Each grey circle represents one FOV. Each solid data point represents the average of all FOVs/mouse (20X magnification). Statistical analysis by unpaired, nonparametric two-tailed Mann-Whitney test. **F)** IHC of Ki67, CC3, and pEGFR in FC1199 orthotopic tumors. Bar graphs represent the mean, and error bars represent the standard deviation. Each grey circle represents one FOV. Each solid data point represents the average of all FOVs/mouse (20X magnification). Statistical analysis by unpaired, nonparametric two-tailed Mann-Whitney test. **G)** IHC of FoxP3 (Tregs) in FC1199 orthotopic tumors. Bar graphs represent the mean, and error bars represent the standard deviation. Each grey circle represents one FOV. Each solid data point represents the average of all FOVs/mouse (20X magnification). Statistical analysis by unpaired, nonparametric two-tailed Mann-Whitney test. **H)** Cd8a (Cd8^+^ T cells) in FC1199 orthotopic tumors. Bar graphs represent the mean, and error bars represent the standard deviation. Each grey circle represents one FOV. Each solid data point represents the average of all FOVs/mouse (20X magnification). Statistical analysis by unpaired, nonparametric two-tailed Mann-Whitney test. **I)** Tumor weight comparison between FC1245 Scr and Fbln3 KD tumors in NSG mice at 5 weeks post-injection. Statistical analysis by unpaired, two-tailed t test. **J)** Gross tumor images collected 5 weeks post-injection of FC1245 Scr and Fbln3 KD and 7 weeks post-injection of FC1199 Scr and Fbln3 KD in C57BL/6 mice. **K)** Gross tumor images collected 5 weeks post-injection of FC1245 Scr and Fbln3 KD in NSG immunodeficient mice.

**Table S1.**
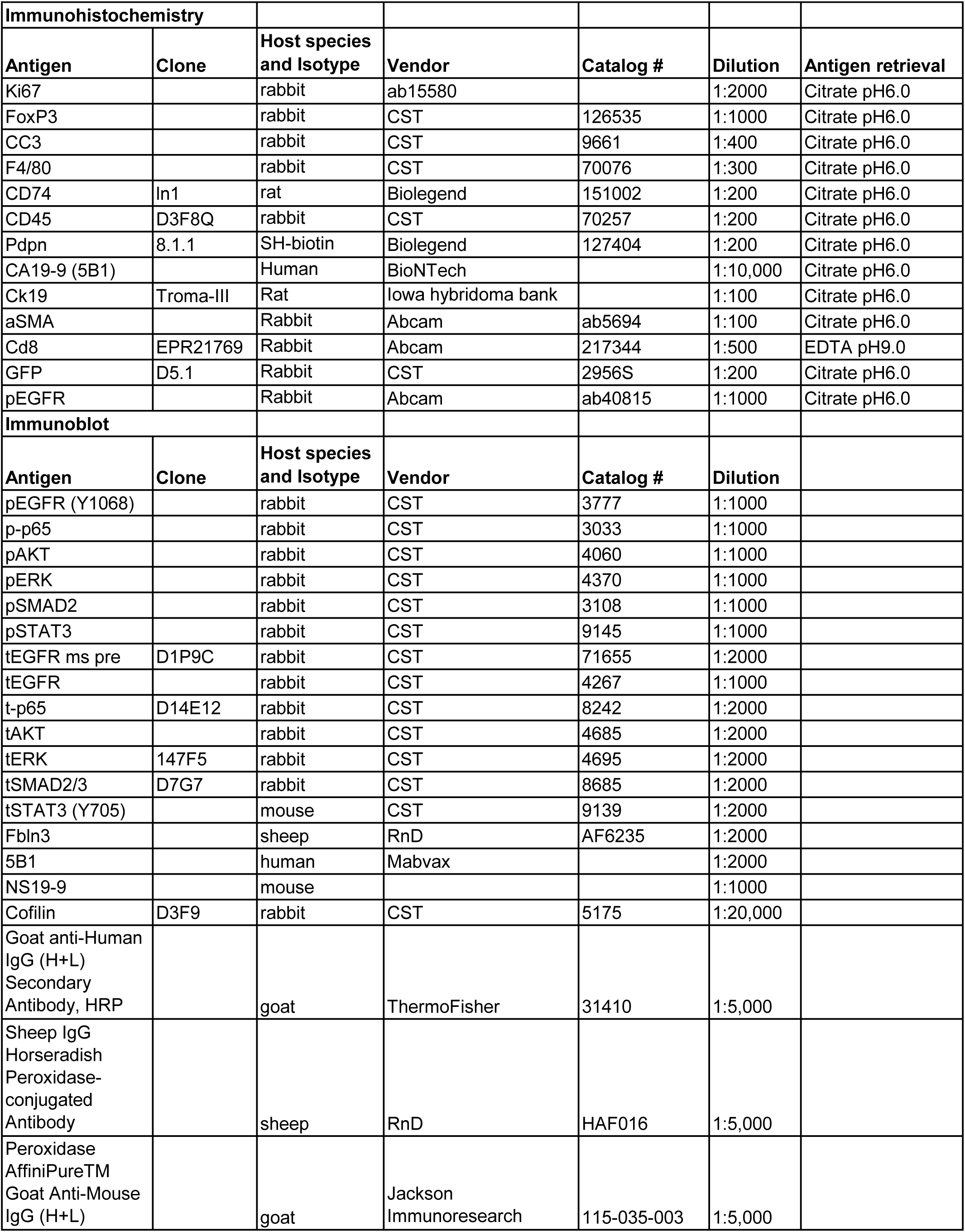
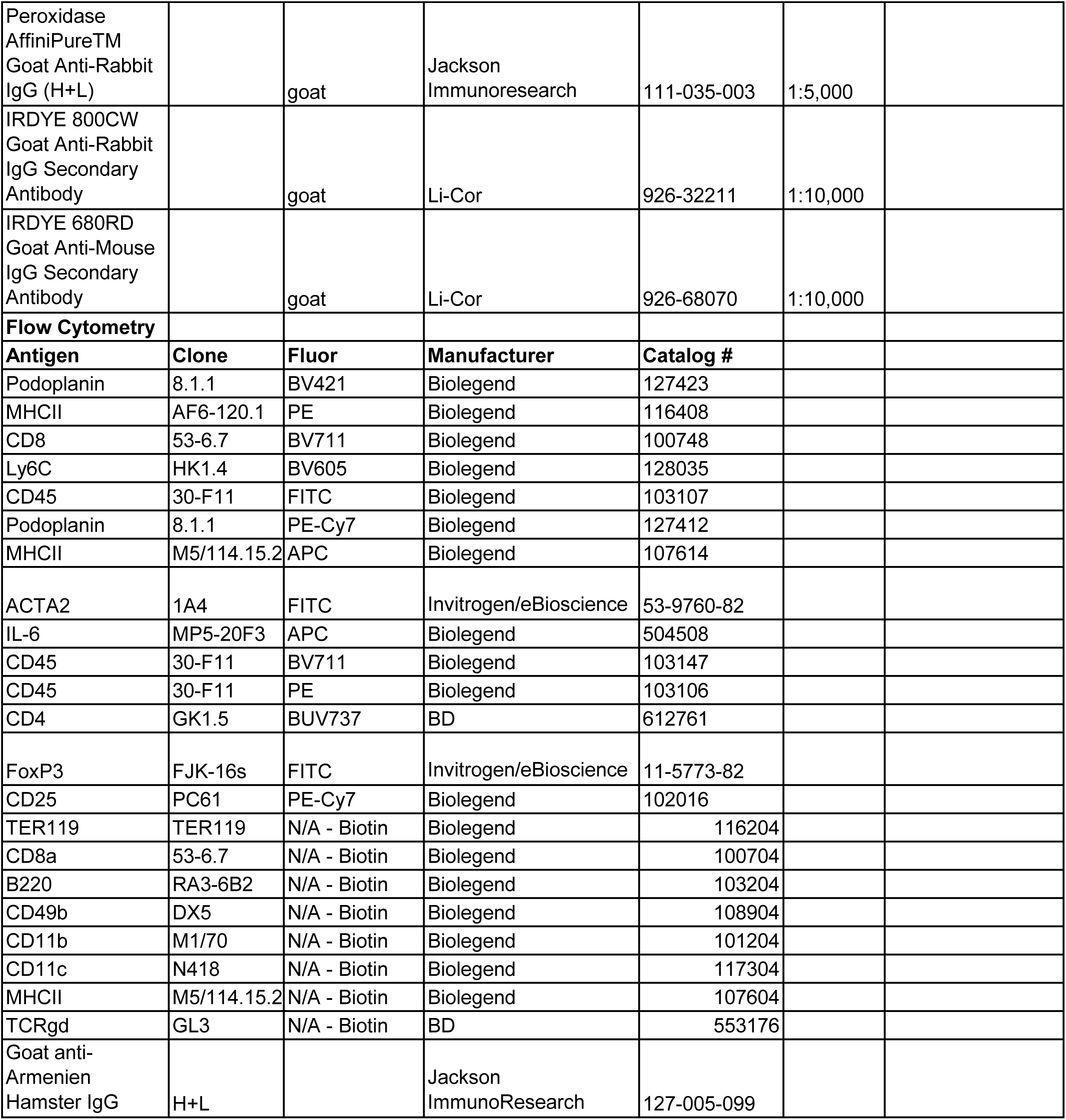

**Table S2.**
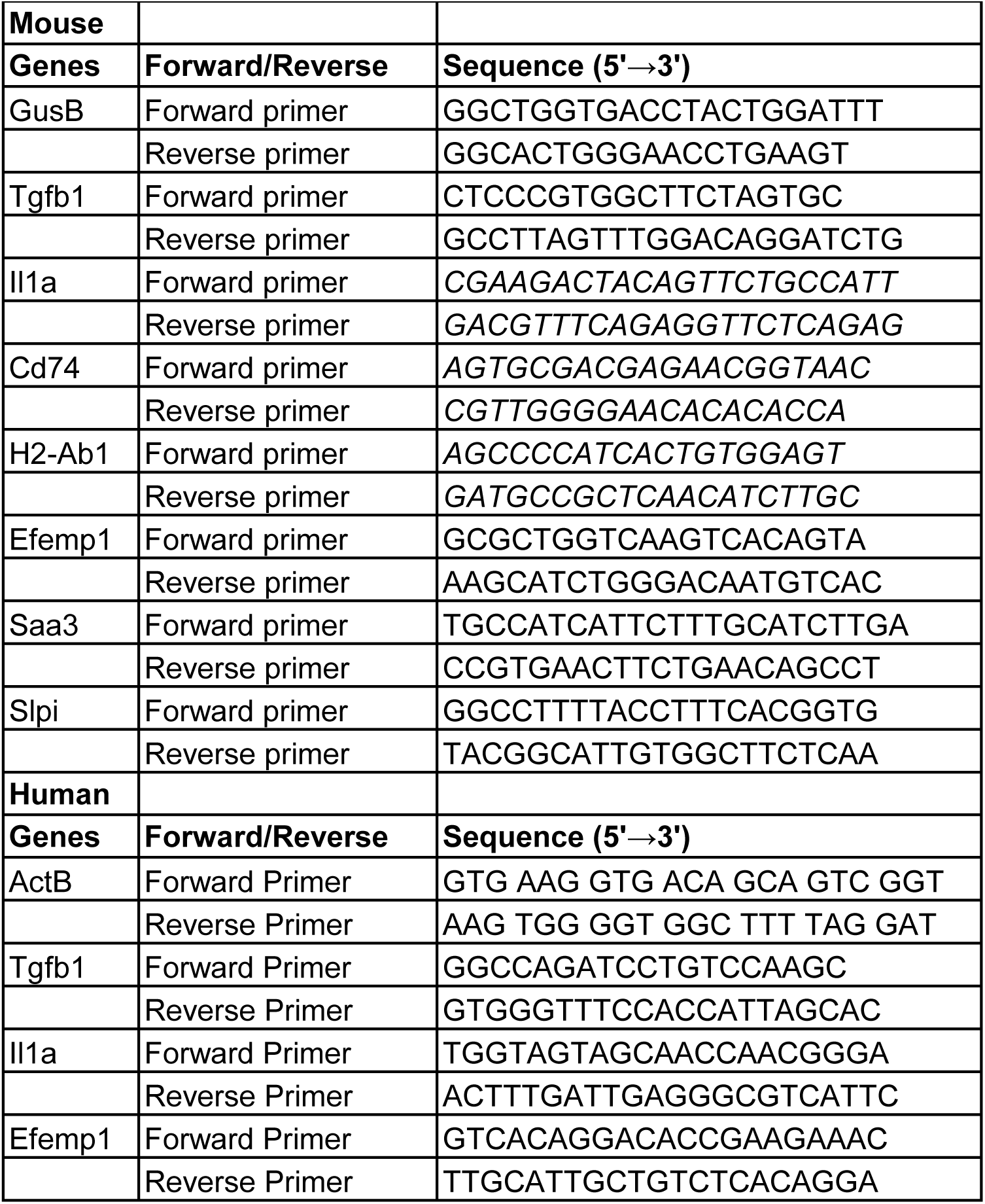

